# A tropane-based ibogaine analog rescues folding-deficient SERT and DAT

**DOI:** 10.1101/2020.07.14.202325

**Authors:** Shreyas Bhat, Daryl A. Guthrie, Ameya Kasture, Ali El-Kasaby, Jianjing Cao, Alessandro Bonifazi, Therese Ku, JoLynn B. Giancola, Thomas Hummel, Michael Freissmuth, Amy Hauck Newman

## Abstract

Missense mutations that give rise to protein misfolding are rare, but collectively, defective protein folding diseases are consequential. Folding deficiencies are amenable to pharmacological correction (pharmacochaperoning), but the underlying mechanisms remain enigmatic. Ibogaine and its active metabolite noribogaine correct folding defects in the dopamine transporter (DAT), but they rescue only a very limited number of folding-deficient DAT mutants, which give rise to infantile Parkinsonism and dystonia. Herein, a series of analogs was generated by reconfiguring the complex ibogaine ring system and exploring the structural requirements for binding to wild type transporters, and for rescuing two equivalent synthetic folding-deficient mutants, SERT-PG^601,602^AA and DAT-PG^584,585^AA. The most active tropane-based analog (**9b**) was also an effective pharmacochaperone *in vivo*, in Drosophila harboring DAT-PG^584,585^AA and rescued six out of 13 disease-associated human DAT mutants *in vitro.* Hence, a novel lead pharmacochaperone has been identified that demonstrates medication development potential for patients harboring DAT mutants.

## Introduction

Ibogaine, one of many alkaloids first isolated from the shrub *Tabernanthe iboga* in 1901 *(Dybowski and Landrin, 1901),* has three interesting pharmacological properties: (i) it is hallucinogenic; presumably the reason why it has been used for centuries by West African tribes in rites of passage *(Wasko et al., 2018; Corkery, 2018).* (ii) It has been reported to mitigate substance use disorders *(Corkery, 2018; Brown and Alper, 2018; Noller et al., 2018).* These actions have also been recapitulated in experimental paradigms, where animals are given the opportunity to self-administer or express their preference for morphine and psychostimulants *(Glick et al., 1991; Glick et al., 1994; Blackburn and Szumlinski, 1997);* (iii) Ibogaine and its principal, active metabolite noribogaine (12-hydroxyibogamine; ref. *Mash et al., 1995)* bind to the transporters for the monoamines serotonin (SERT), dopamine (DAT) and norepinephrine (NET) *(Mash et al., 1995; Sweetnam et al., 1995).* Moreover, ibogaine and noribogaine were the first compounds that were shown to rescue folding-deficient versions of SERT *(El-Kasaby et al., 2010; El-Kasaby et al., 2014; Koban et al., 2015)* and of DAT *(Kasture et al., 2016; Beerepoot et al., 2016; Asjad et al., 2017).* This pharmacochaperoning action of ibogaine and noribogaine is of therapeutic interest because folding-deficient mutants of DAT give rise to the dopamine transporter deficiency syndrome or DTDS *(Kurian et al., 2011; Ng et al., 2014).* DTDS is a hyperkinetic movement disorder in DAT deficient patients that progresses into Parkinsonism and dystonia. This disease manifests generally in the first 6 months post birth (infantile) or occasionally during childhood, adolescence or adulthood (juvenile). The cell surface levels of DAT variants associated with DTDS are typically non-detectable (in infantile cases) or severely reduced (in juvenile cases) due to their misfolding and subsequent retention within the endoplasmic reticulum (ER). While the use of ibogaine or noribogaine as a medication for DTDS is warranted, its hallucinogenic profile will likely preclude its therapeutic usefulness in this patient population (*Wasko et al., 2018*).

The closely related monoamine transporters SERT (*SLC6A4*), DAT *(SLC6A3)* and NET *(SLC6A2)* form a branch of the solute carrier-6 (SLC6) family of secondary active transporters (*Kristensen et al., 2011*). They modulate monoaminergic neurotransmission by retrieving their eponymous substrates from the synaptic cleft, which supports replenishing of vesicular stores. SERT and - to a lesser extent - NET are the most important targets for antidepressants, which act as inhibitors. For example, SSRIs (selective serotonin reuptake inhibitors) are used to treat major depression, obsessive-compulsive disorders, and general anxiety disorders. The therapeutic indication for DAT inhibition is more restricted (e.g., methylphenidate for attention-deficit-hyperactivity disorder or modafinil for sleep disorders). However, DAT is a prominent target for illicit drugs (e.g., cocaine and amphetamines). This is also true to some extent for SERT, which is the target of 3,4-methylenedioxymethamphetamine (MDMA, ecstasy) and its congeners. Thus, the chemical space for these transporter targets has been explored *(Sitte and Freissmuth, 2015).* Of note, monoamine transporter ligands can range from full substrates to typical inhibitors, as well as atypical inhibitors, depending on chemical structure and transporter conformation *(Schmitt et al., 2013; Reith et al., 2015; Bhat et al., 2019)* Typical inhibitors (e.g. cocaine, most antidepressants) bind to and trap the transporter in the outward facing state thus precluding any subsequent conformational transition required for entry of the protein into a transport mode. In contrast, the substrate-bound transporter enters an occluded state. Substrate translocation is initiated by opening of an inner gate, which releases the substrate together with co-substrate ions (Na^+^ and Cl^-^) on the intracellular side (*Sitte and Freissmuth, 2015*). Full substrates allow the transporter to undergo its transport cycle in a manner indistinguishable from cognate neurotransmitter. Full substrates, which differ from neurotransmitters in their cooperative interaction with the co-transported sodium, act as amphetamine-like releasers by driving the transporter into an exchange mode (*Hasenhuetl et al., 2019)* Partial substrates/releasers support the transport cycle/exchange mode albeit less efficiently than full substrates/releasers, because they bind tightly to conformational intermediates and thus preclude rapid transitions *(Bhat et al., 2017).* Atypical inhibitors trap the transporter in conformations other than the outward facing state. Ibogaine is an atypical inhibitor that binds the inward-facing state of SERT *(Jacobs et al., 2007; Bulling et al., 2012; Burtscher et al., 2018; Coleman et al., 2019)* and, presumably, of DAT and NET. Several arguments support the conjecture that the folding trajectory of monoamine transporters proceeds through the inward facing state *(Freissmuth et al., 2017),* thus providing a mechanistic basis for rationalizing the pharmacochaperoning action of ibogaine (*El-Kasaby et al., 2010; El-Kasaby et al., 2014; Koban et al., 2015; Kasture et al., 2016; Beerepoot et al., 2016; Asjad et al., 2017)* and of other compounds *(Bhat et al., 2017).*

Ibogaine and its metabolite noribogaine are the most efficacious pharmacochaperones for folding-deficient versions of SERT and DAT identified to date and provide templates to generate promising new leads. In this study, we explored the chemistry of ibogaine to broaden the efficacy profile for this drug in rescuing misfolded SERT and DAT mutants. Ibogaine has a complex and rigid structure, which until recently has been largely unexplored *(Kruegel et al., 2015, Gassaway et al., 2016).* Here, we generated a series of novel ibogaine analogs by investigating the chemical space surrounding the parent molecule using a two-pronged approach: 1) Deconstructing ibogaine by introducing flexible hydrocarbon linkers that connect the indole ring to either a isoquinuclidine ring or a tropane ring and 2) Reconfiguring and completely substituting the isoquinuclidine ring of ibogaine with the tropane ring system. This allowed for defining structural determinants required for high-affinity binding to wild type SERT and DAT and for pharmacochaperoning two synthetic folding-deficient mutants: SERT-PG^601,602^AA and the orthologous DAT-PG^584,585^AA. Based on these experiments, we identified a novel fluorinated tropane-based analog, which was more potent and more efficacious than the parent compound in rescuing misfolded versions of SERT and DAT, including disease-relevant DTDS mutants. Importantly, this compound was active *in vivo* and restored sleep to flies harboring a misfolded DAT mutant.

## Results

### Synthesis

Ibogaine can be viewed as a serotonin analog, where the basic nitrogen is fixed in space by a fused bicyclic ring structure *(cf.* **Table 1**). Two strategic approaches were undertaken that yielded a number of unique products to the ibogaine series in relatively few synthetic steps. The first approach (**Scheme 1A**) involved a deconstructive strategy, whereby the indole ring was disconnected from the isoquinuclidine ring structure, thereby offering greater flexibility along with a comparable number of hydrocarbon atoms by either retaining the isoquinuclidine ring (**4a–c**) or replacing it with a tropane ring (**3a–f**). The second approach (**Scheme 1B**) aimed to reconfigure and completely substitute the fused isoquinuclidine ring of ibogaine with the tropane ring system conferring the novel intermediate amides (**8a–c**) and their reduced tertiary amine analogs (**9a–d**). Of note, the tropane ring is a hallmark structure in many classical monoamine transporter inhibitors (e.g., cocaine, benztropine) and thus was envisioned to potentially retain binding affinities at DAT and SERT. In both approaches, the ethyl group of ibogaine was eliminated to save on synthetic effort. In addition, we surmised that the ethyl group was immaterial for binding affinity (see below).

**Table 1:** Radioligand uptake and binding data at DAT and SERT^a^

|  | Uptake inhibition |  | Binding inhibition |  | ratio |  | Pharmacochaperoning |  |  |  |
| --- | --- | --- | --- | --- | --- | --- | --- | --- | --- | --- |
| | $IC_{50} \pm S.D.$ | | $K_i \pm S.D.$ | | $IC_{50}/K_i$ | | $EC_{50} \pm S.D.$ | $V_{max}$ | $EC_{50} \pm S.D.$ | $V_{max}$ |
| | $\mu M$ | | $\mu M$ | | | | $\mu M$ | % | $\mu M$ | % |
|  | WT-SERT | WT-DAT | WT-SERT | WT-DAT | WT-SERT | WT-DAT | SERT-PG<sup>601,602</sup> AA |  | DAT-PG<sup>584,585</sup> AA |  |
| Ibogaine | 8.2 ± 3.5 | 22.1 ± 6.3 | 7.4 ± 0.7 | 5.3 ± 1.7 | 1.1 | 4.2 | 2.6 ± 1.9 | 103 ± 16 | 19.6 ± 4.9 | 88 ± 16 |
| Noribogaine | 1.2 ± 0.5 | 15.5 ± 7.8 | 0.6 ± 0.1 | 4.0 ± 1.8 | 2.1 | 3.9 | 4.6 ± 1.2 | 100 ± 8 | 17.1 ± 6.1 | 100 ± 13 |
| 3a | 9.1 ± 3.1 | 25.6 ± 10.2 | 1.2 ± 0.2 | 3.1 ± 0.9 | 7.8 | 8.2 | NC* | 19 ± 6* | NC* | 15 ± 10* |
| 3b | 6.5 ± 5.2 | 24.4 ± 17.9 | 1.85 ± 0.3 | 1.7 ± 0.1 | 3.5 | 14.4 | 12.9 ± 6.3 | 60 ± 17 | NC* | 15 ± 8* |
| 3c | 0.5 ± 0.1 | 9.09 ± 1.64 | 0.19 ± 0.03 | 4.4 ± 1.4 | 2.4 | 2.1 | 2.7 ± 0.78 | 57 ± 13 | 17.1 ± 5.6 | 47 ± 18 |
| 3d | 11.9 ± 1.6 | 58.2 ± 41.4 | 4.5 ± 0.4 | 8.8 ± 4.94 | 2.6 | 6.6 | NC* | 11 ± 3* | NC* | 20 ± 15* |
| 3e | 21.6 ± 17.4 | 179 ± 94 | 13.0 ± 2.7 | 10.2 ± 3.8 | 1.7 | 17.6 | NC* | 38 ± 19* | NC* | 15 ± 9* |
| 3f | 0.2 ± 0.1 | 78.0 ± 27.9 | 0.12 ± 0.02 | 21.8 ± 9.4 | 1.6 | 3.6 | NC* | 8 ± 2* | NC* | 13 ± 10* |
| 4a | 4.6 ± 2.63 | 6.5 ± 2.1 | 0.81 ± 0.03 | 1.9 ± 0.2 | 5.7 | 3.5 | NC* | 25 ± 6* | NC* | 15 ± 9* |
| 4b | 8.2 ± 1.0 | 11.9 ± 1.6 | 1.7 ± 0.3 | 1.5 ± 0.3 | 4.8 | 7.7 | NC* | 29 ± 8* | NC* | 10 ± 6* |
| 4c | 0.3 ± 0.2 | 11.8 ± 9.1 | 0.13 ± 0.34 | 2.6 ± 0.7 | 2.5 | 4.5 | 0.7 ± 0.3 | 68 ± 16 | NC* | 24 ± 8* |
| 8a | ND | ND | >100 | 39.7 ± 27.9 | ND | ND | ND | ND | ND | ND |
| 8b | ND | ND | >100 | 51.2 ± 14.6 | ND | ND | ND | ND | ND | ND |
| 8c | ND | ND | >100 | 45.78 ± 6.5 | ND | ND | ND | ND | ND | ND |
| 9a | 6.9 ± 3.9 | 16.7 ± 11.2 | 4.4 ± 0.2 | 3.6 ± 0.2 | 1.6 | 4.6 | 3.05 ± 2.3 | 94 ± 23 | 18.9 ± 2.9 | 51 ± 21 |
| 9b | 0.4 ± 0.2 | 7.2 ± 3.8 | 0.18 ± 0.01 | 0.8 ± 0.3 | 2.3 | 8.9 | 0.06 ± 0.04 | 100 ± 6 | 25.7 ± 6.3 | 673 ± 245 |
| 9c | 29.9 ± 16.4 | 10.8 ± 9.3 | 7.3 ± 0.5 | 3.1 ± 0.3 | 4.1 | 3.5 | 2.1 ± 0.8 | 81 ± 20 | 19.2 ± 8.7 | 53 ± 13 |
| 9d | 0.9 ± 0.5 | 15.2 ± 9.7 | 0.78 ± 0.04 | 2.9 ± 0.3 | 1.2 | 5.3 | 1.4 ± 0.8 | 89 ± 19 | 5.8 ± 1.9 | 49 ± 30 |

<sup>a</sup>Each $IC_{50}$ , $K_i$ , $K_m$ and $V_{max}$ value represent data from at least 3 independent experiments, each performed in triplicate. $V_{max}$ refers to the maximum transport restored by noribogaine, which was set at 100% to account for inter-assay variation. Experimental procedures are described in detail in Methods. NC\*: not calculable: a reliable $EC_{50}$ could not be estimated, because functional rescue was too low; in this instance $V_{max}$ \* refers to the activity restored by 30 $\mu M$ of the compound. ND: not determined.

**Scheme 1.**
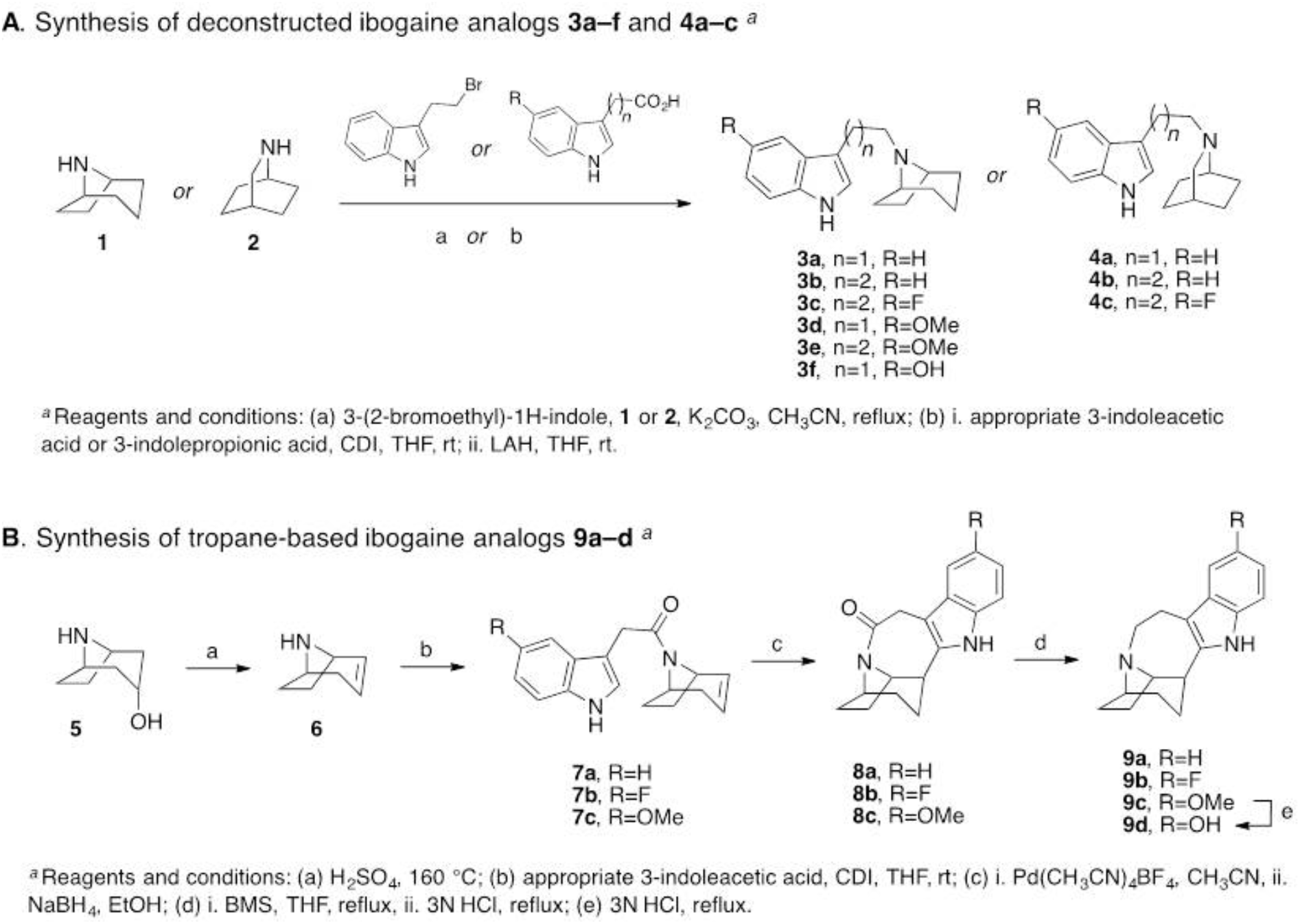
Synthetic scheme of the ibogaine analogs

The synthetic schemes of the two approaches are shown in **Scheme 1**. 8-Azabicyclo[3.2.1]octane (nortropane, (**1**)) or 2-azabicyclo[2.2.2]octane (**2**) was directly linked at the *N*-position with various indole substituents via either nucleophilic substitution with 3-(2-bromoethyl)-1H-indole or by tandom 1,1’-carbonyldiimidazole (CDI) coupling with an appropriate 3-indoleacetic acid or 3-indolepropanoic acid followed by lithium aluminum hydride (LAH) reduction to give the desired products (**3a–f**, **4a–c**) as free bases (**Scheme 1A**). The second approach (**Scheme 1B**) started similarly following the dehydration of nortropine (**5**) in sulfuric acid to give racemic nortropidene (**6**), which was linked with an appropriate 3-indoleacetic acid substituent via CDI coupling. Next, similarly to Trost *et al.* (*Trost et al.,1978)* and Kruegel *et al. (Kruegel et al., 2015)* in their syntheses of ibogamine, electrophilic palladium promoted cyclization of racemic **7a–c**, followed by sodium borohydride (NaBH_4_) reduction, gave the racemic products **8a–c**. This key step produced the corresponding amides in good yields (63-81% yields, see Experimental Methods in S. I. for details), without necessitating chromatographic separation. Notably, Trost et al. (*Trost et al.,1978)* and Kruegal et al. *(Kruegel et al., 2015)* reported much lower isolated yields of ibogamine (19% and 33%, respectively) with this chemistry, by using the isoquinuclidine core and corresponding amine instead. Finally, the amide reduction of racemic **8a**–**c** was accomplished with borane dimethyl sulfide complex (BMS) to give racemic **9a–d**, where **9d** was formed from the *O*-demethylation of **9c** following extended stirring at reflux, in 3N HCl.

### Structure activity-relationship of ibogaine analogs at SERT and DAT

Ligand affinities at DAT and SERT can be determined by measuring their ability to displace a radioligand from the transporter or to block substrate uptake. During substrate translocation, transporters undergo a conformational cycle. Radioligands are high-affinity inhibitors, which bind to the outward facing state. In addition, both the membrane potential and the asymmetric ion distribution are absent in binding assays with membrane preparations. Hence, affinity estimates for some ligands can differ substantially, if inhibition of substrate uptake and displacement of substrate are assessed. Because of conformational trapping in the absence of an ion gradient, these differences can reach several orders of magnitudes (*Bhat et al., 2017*). Accordingly, we determined the affinity of our novel analogs by measuring their ability to both, displace a radioligand in membranes prepared from rat brain stem (**Supplementary Fig 1**, SERT) and rat striatum (**Supplementary Fig 2**, DAT) and inhibit cellular uptake in transfected cells (**Supplementary Figs 3 and 4** for SERT and DAT, respectively). Noribogaine is known to bind SERT with higher affinity, in both binding and uptake inhibition assays, over the methoxylated parent compound, ibogaine, and our data confirm this in **Table 1**. However, as previously reported *(Mash et al., 1995),* ibogaine and noribogaine have similar affinities for DAT (also seen in **Table 1**). It is evident from the data in **Table 1** that the *C*-12 substitutions primarily determine affinity to SERT in all analogs tested. Methoxy-(**3d**, **3e**, **9c**) or hydroxy-substitution (**3f**, **9d**) in this 12-position has little effect on DAT affinity as compared to the unsubstituted analogs (**3a**, **3b**, **4a**, **4b**, **8a**, **9a**), but the hydroxy-substituted analogs have higher affinity at SERT (**3f**, **9d**). Interestingly, when the hydroxyl group is substituted with fluorine, compounds **3c**, **4c** and **9b** also exhibit similar affinities at SERT to the hydroxy-analogs. Only **9b** showed submicromolar affinity for DAT, suggesting a potentially pivotal role for fluorine substitution. In addition, compounds **9c** and **9d** are ibogaine and noribogaine analogs, respectively, in which the 2-ethyl group has been removed and the isoquinuclidine core has been replaced with a tropane ring. The affinities of these compound pairs were similar at DAT and SERT (**9c**/ibogaine v. **9d**/noribogaine). These data supported our prediction that the ethyl-group on the isoquinuclidine ring of the ibogamine structure is dispensable and that the isoquinuclidine core can be replaced by tropane. The rigidity imparted by the azepine-ring (compounds **9a–d**, ibogaine, noribogaine) is a minor determinant of affinity. Indeed, the affinity of compounds **3c** and **9b**, for instance, were comparable.

The amide analogs, in which the basic amine was neutralized, had substantially reduced affinities (*cf.* compounds **8a–c** and **9a–c** in **Table 1**) demonstrating the necessity for a protonatable nitrogen. In general, the affinities obtained from binding displacement were higher than those estimated from uptake inhibition. Despite this difference, in the two affinity estimates in SERT, the rank order of potency determined by uptake inhibition was reasonably similar to that determined by binding: analogs with hydroxy- and fluorine substituents were more potent in blocking substrate uptake by SERT than the methoxy- and unsubstituted counterparts (**Table 1**).

In DAT, all fluorinated analogs (**3c**, **4c** and **9b**) also ranked among the most potent blockers of DAT uptake (**Table 1**). However, affinity estimates from binding and uptake inhibition differed in part substantially with IC_50_/K_i_ ratios ranging from 2 to 14 (**Table 1**). Variations in the ratio of IC_50_/K_i_ (see **Table 1**) are indicative of differences in conformational trapping *(Bhat et al., 2017)* rather than of species differences (rat v. human transporters). More importantly, conformational trapping is predictive of a pharmacochaperoning action (*Bhat et al., 2017).* Accordingly, we determined if some of these analogs were capable of rescuing folding-deficient mutants of SERT and DAT.

### Pharmacochaperoning action of the ibogaine analogs on SERT-PG^601,602^AA and on DAT-PG^584,585^AA

Several folding-deficient versions of SERT have been generated and characterized to understand the actions of pharmacochaperones (*El-Kasaby et al., 2010; El-Kasaby et al., 2014; Koban et al., 2015*). The mutant SERT-PG^601,602^AA is stalled at an early stage of the folding trajectory (*El-Kasaby et al. 2014*). Hence, based on its severe phenotype, we selected this mutant to examine and compare the ability of our ibogaine analogs (i.e., compounds **3a– 3f, 4a–4c, 9a–9d**) with ibogaine and noribogaine to act as pharmacochaperones. Misfolded transporters are retained in the ER and thus unavailable to accomplish their eponymous actions, that is cellular uptake of substrate. Pretreatment of cells with a pharmacochaperone restores folding of the protein and thus its subsequent delivery to the cell surface (*El-Kasaby et al., 2010; El-Kasaby et al., 2014; Koban et al., 2015; Kasture et al., 2016; Beerepoot et al., 2016; Asjad et al., 2017).* This can be readily monitored by measuring cellular substrate uptake. As shown in **Fig. 1A**, in cells expressing SERT-PG^601,602^AA, there is negligible uptake of [^3^H]5-HT (<<10%) when compared to those expressing wild type SERT.

**Figure 1.**
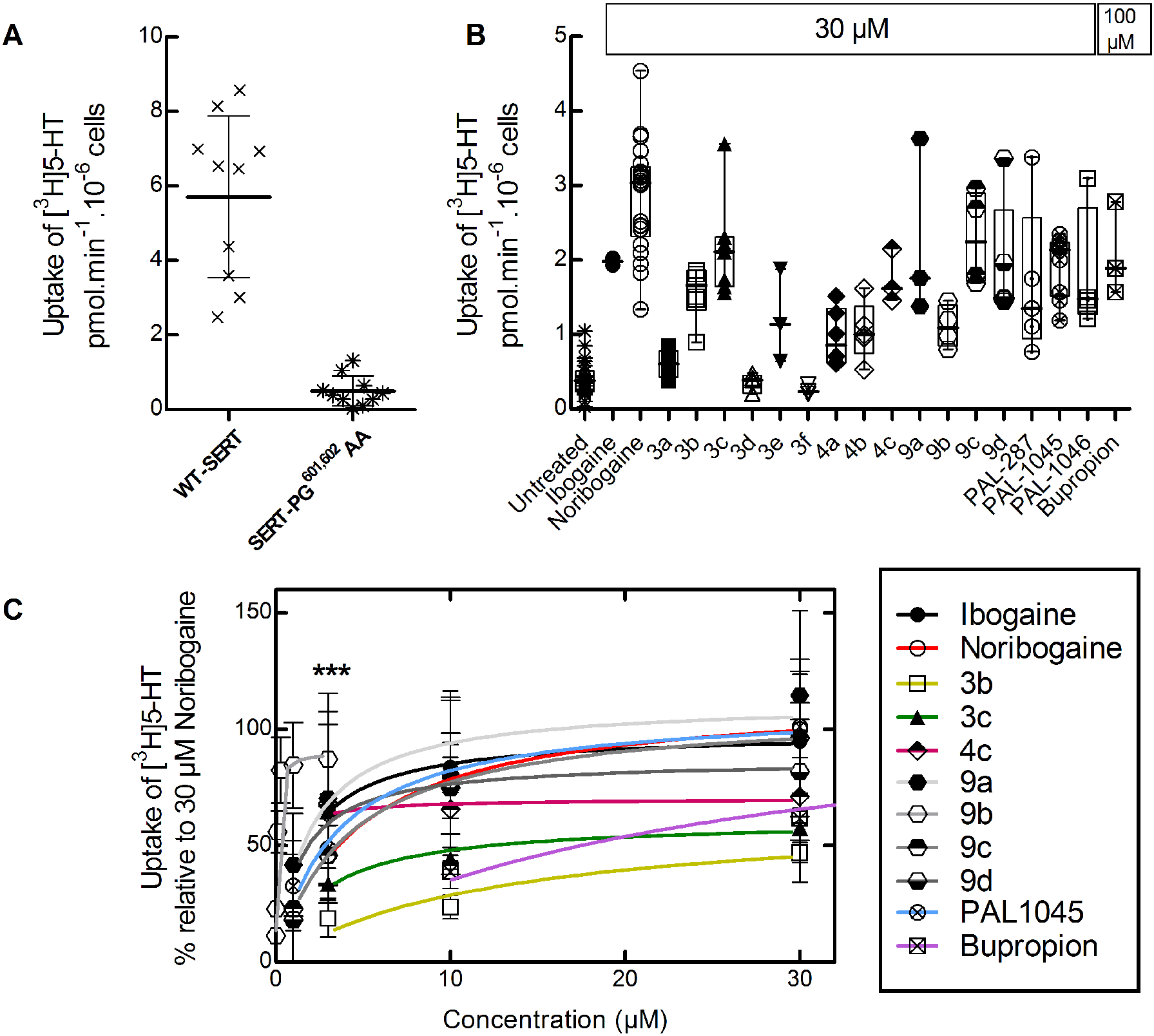
[^3^H]5-HT uptake in HEK293 cells expressing hSERT-PG^601,602^AA after pre-incubation with ibogaine analogs. ***A.*** Comparison of [^3^H]5-HT uptake by HEK293 cells transiently expressing wild type (WT) hSERT or hSERT-PG^601,602^AA. Each symbols represents the result from an individual experiment (done in triplicate); means± S.D. are also shown. ***B.*** hSERT-PG^601,602^AA expressing cells were incubated in the presence of the indicated compounds (30 μM in all instances but bupropion, 100 μM). After 24 h, [^3^H]5-HT uptake was determined as outlined under “Methods”. Values from individual experiments (done in triplicate) are shown as dots; box plots show the median and the interquartile range. ***C.*** Concentration-response curves for pharmacochaperoning by the indicated compounds (selected as positive hits from *panel B).* Rescued uptake was normalized to that achieved by 30 μM noribogaine (=100%) to account for inter-experimental variations. ***, E_max_ for **9b** was achieved at 3 μM, uptake inhibition was observed after pre-incubation with higher concentrations. The solid lines were drawn by fitting the data to the equation for a rectangular hyperbola (for EC_50_ and E_max_ of mutant rescue see **Table 1**). Data were obtained in at least three independent experiments carried out in triplicate. The error bars indicate S.D.

In cells, which were preincubated for 24 h in the presence of 30 μM ibogaine, noribogaine and other pharmacochaperones, i.e., the naphthyl-propanamine series (PAL-287, PAL-1045 and PAL-1046, *Rothman et al., 2012; Bhat et al., 2017)* and 100 μM bupropion *(Beerepoot et al., 2016),* substrate uptake supported by SERT-PG^601,602^AA increased to levels, which corresponded to 25-40% of transport activity of wild type SERT (**Fig. 1B**). Similarly, if the cells were preincubated in the presence of 30 μM of the new ibogaine analogs, appreciable levels of transport were seen with compounds **3b**, **3c**, **4c**, **9a, 9c** and **9d** (**Fig. 1B**). Additional experiments (not shown) confirmed that compound **9b** was more effective at concentrations <10 μM than at 30 μM. Hence, it was also included in the list of positive hits, which were further investigated to determine their potency and their efficacy as pharmacochaperones. We compared their action in individual transient transfections by normalizing their pharmaco-chaperoning activity to the transport activity restored by 30 μM noribogaine (**Fig. 1C**). Known pharmacochaperones (PAL-1045, bupropion, ibogaine and noribogaine) were also examined as reference compounds. All compounds belonging to the **9a–9d** series were as effective as noribogaine, but compound **9b** had an EC50 of ~60 nM (curve represented as *** in **Fig. 1C**). The other fluoro-analog, **4c,** had an EC50 of ~600 nM but was less efficacious (E_max_ ~70% of that of noribogaine). Other compounds, including noribogaine (EC_50_ = ~3 μM), rescued SERT-PG^601,602^AA with EC_50_-values in the low to mid μM range (**Table 1**).

Our data suggests **9b** to be the most potent of all compounds tested in rescuing SERT-PG^601,602^AA. We visualized the glycosylation state of SERT-PG^601,602^AA by immunoblotting to confirm the pharmacochaperoning action of **9b** by an independent approach. During their synthesis in the ER, membrane proteins undergo N-linked core glycosylation; this core glycan, which can be removed by endoglycosidase H, is thus present on ER-retained misfolded mutants. Rescued mutants are exported to the Golgi, where they acquire additional sugar moieties. The resulting complex glycan structure is resistant to cleavage by endoglycosidase H. The core glycosylated protein is homogeneous and smaller in size than the mature glycosylated version, which is heterogeneous due to the stochastic nature of complex glycosylation. Accordingly, in lysates prepared from transiently transfected cells, wild type SERT was visualized by immunoblotting as a band migrating at 75 kDa and broad smear coalescing from a collection of bands in the range of 90 to 110 kDa (left hand lane, **Fig. 2A**). The size of the lower band was reduced after cleavage by endoglycosidase H (*cf.* first and second lane in **Fig. 2C**) confirming that it corresponded to the core glycosylated band (“C”). In contrast, the migration of the collection of upper bands was insensitive to cleavage by endoglycosidase H confirming that they had acquired the mature glycan *(cf.* bands “M” in first and second lane in **Fig. 2C**). In contrast, lysates from cells expressing SERT-PG^601,602^AA contain predominantly the core glycosylated protein (second lane in **Fig. 2A** showing control cells). As predicted from its ability to restore substrate uptake in pretreated cells (**Fig. 1B,C**), **9b** induced the appearance of slowly migrating forms of SERT-PG^601,602^AA in lysates prepared after preincubation of the cells in a concentration-dependent manner (lanes 3 to 8 in **Fig. 2A**). Plotting this concentration response revealed an EC_50_ of 134 ± 30 nM (**Fig. 2B**), which is in excellent agreement with that calculated from restoration of substrate uptake (**Fig. 1C** and **Table 1**). We confirmed that the mature glycosylated species of the rescued mutant is resistant to cleavage by endoglycosidase H (shown for lysates from cells compound **9b**-pretreated cells in **Fig. 2C**).

**Figure 2.**
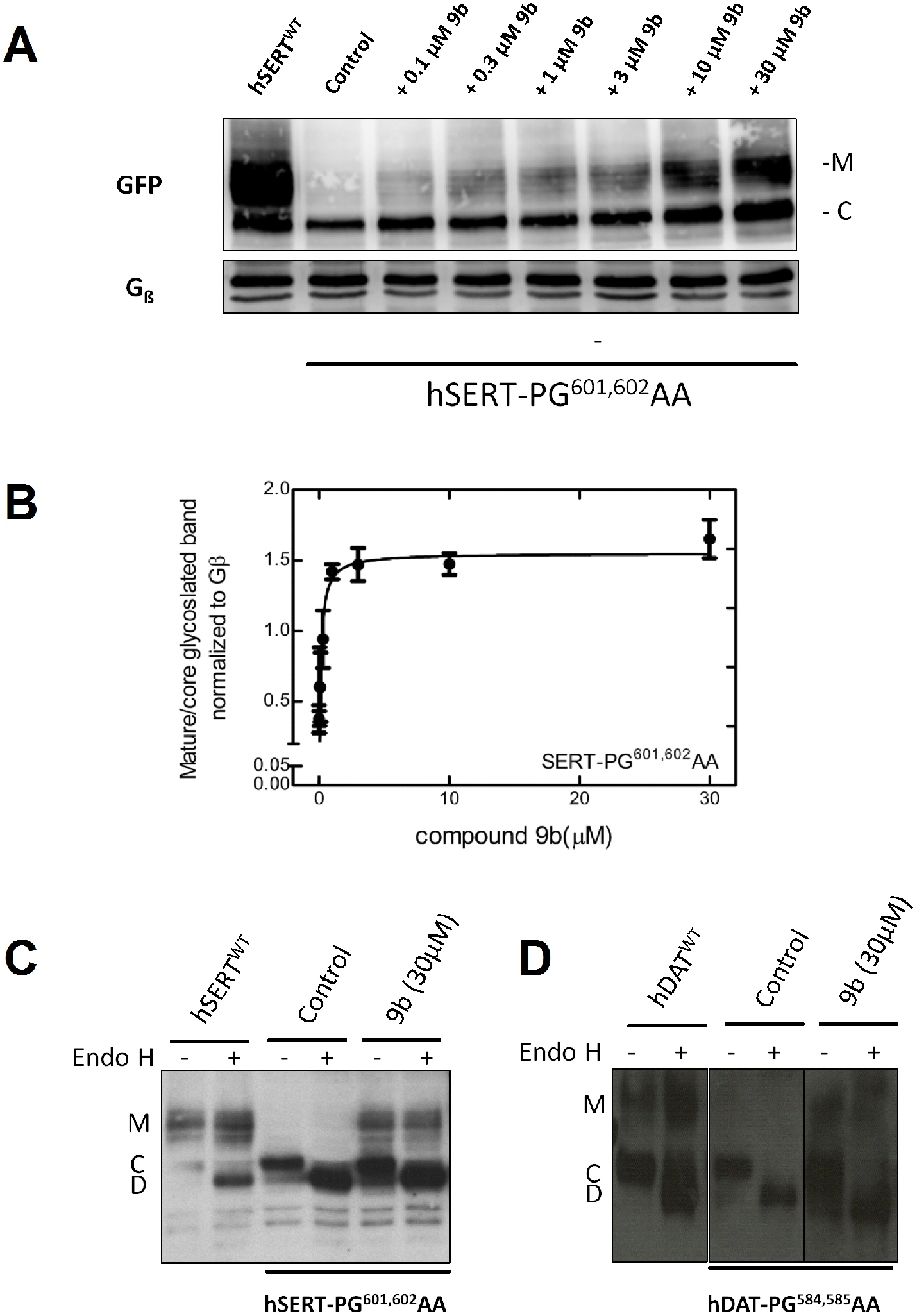
Enhanced mature glycosylation of hSERT-PG^601,602^AA or hDAT-PG^584,585^AA in cells pre-incubated with 9b. ***A.*** HEK293 cells transiently expressing hSERT-PG^601,602^AA were incubated in the absence (negative control = lane 2) or presence of the indicated concentrations of **9b** for 24 h. Cells expressing wild type (WT) hSERT were the positive control (first labeled as hSERT^WT^). Membrane proteins extracted from these cells were denatured, electrophoretically resolved and transferred onto nitrocellulose membranes. The blots were incubated overnight at 4 °C with anti-GFP (top) or anti-Gβ (bottom, loading control) antibodies. The immunoreactive bands were detected with using a horseradish peroxidase-conjugated secondary antibody. ***B.*** Concentration-response curve generated from three independent experiments carried out as in *A.* The ratio of mature (M) to core glycosylated band (C) was quantified densitometrically, normalized to the density of Gβ (loading control) and compared with the ratio observed in each blot for untreated control cells. The density ratios for untreated control cells and those exposed to 30μM compound **9b** (30 μM) were 0.38 + 0.09 and 1.65 + 0.14, respectively. The solid line was drawn by fitting the data to the equation for a rectangular hyperbola. The EC_50_ of rescue was 134 + 30 nM. ***C.*** and ***D.*** In separate experiments, lysates were prepared from cells expressing the SERT-PG^601,602^AA and DAT-PG^584,585^AA mutants treated in the absence or presence of 30 μM **9b and** subjected to enzymatic digestion by endoglycosidase H (Endo H). Endo H specifically cleaves core glycans (C) to generate lower molecular weight deglycosylated fragments (D). Mature glycosylated bands (M) are resistant to the actions of Endo H.

We posited that the potency and efficacy of the ibogaine analogs in pharmacochaperoning SERT and DAT would best be compared in the equivalent folding-deficient mutants. The residues P^601^ and G^602^ in SERT are conserved in DAT at positions P^584^ and G^585^, respectively. Thus, we created DAT-PG^584,585^ AA and verified that it was retained within the cell (not shown), accumulated predominantly as core glycosylated species (*cf.* lanes 3 & 4 and lanes 1 & 2 in **Fig. 2D**) and only mediated very low level of substrate uptake **(Fig. 3A**). The folding deficiency of this mutant is not only expected based on the homology of SERT and DAT but also predicted from earlier work: mutation of glycine^585^ to an alanine in DAT suffices to completely abrogate surface expression of the transporter *(Miranda et al., 2004).* As shown in **Fig. 3B**, if cells transiently expressing DAT-PG^584,585^AA were preincubated in the presence of 30 μM noribogaine, there was a robust increase (~6 fold) in substrate uptake. Of the other known pharmacochaperones, ibogaine, PAL-287 and PAL-1046 (but not PAL-1045) were also active. Of the ibogaine analogs, compounds **3c**, **4c** (the fluorinated open-ring analogs) and **9a, 9c** and **9d** rescued the function of DAT-PG^584,585^AA to a modest extent. In contrast, compound **9b** was a very efficacious pharmacochaperone for DAT-PG^584,585^AA mutant with transport activity exceeding that restored by noribogaine by ~4-fold. Thus, after preincubation in the presence of compound **9b**, cells expressing DAT-PG^584,585^AA recovered up to ~40% of the uptake velocity seen in cells expressing wild type DAT (*cf.* **Figs. 3A** & **3B**) and mature glycosylated protein accumulated to appreciable levels (lanes 5 and 6 in **Fig. 2D**). We selected compounds **3c**, **4c**, **9a**–**9d**, PAL-287 and PAL-1046 as positive hits. Cells transiently expressing DAT-PG^584,585^AA were preincubated with increasing concentrations of these compounds and substrate uptake was subsequently determined to obtain concentrationresponse curves for their pharmacochaperoning activity. When compared to the reference compounds ibogaine and noribogaine, compounds **3c**, **4c**, **9a**, **9c**, **9d**, PAL-287 and PAL-1046 were all less efficacious, and the EC50-values of compounds **3c**, **4c**, **9a**, **9d** and PAL-1046 were lower than those of ibogaine and noribogaine (**Fig. 3C**, upper panel). As predicted from the screening assays summarized in **Fig. 3B**, compound **9b** was substantially more efficacious than noribogaine or ibogaine, but the EC_50_-values of these three compounds were comparable (**Fig. 3C**, lower panel; **Table 1**).

**Figure 3.**
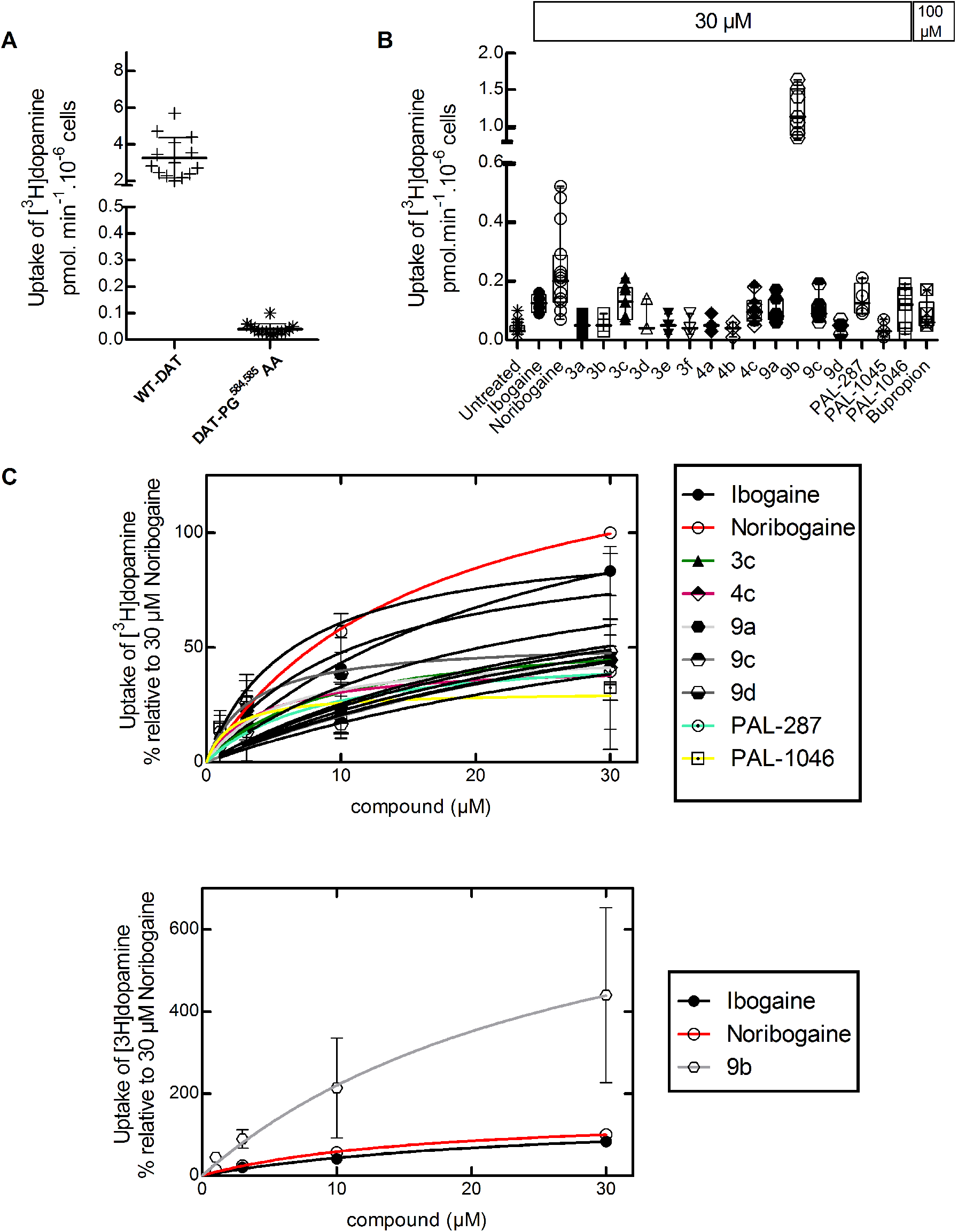
[^3^H]Dopamine uptake in HEK293 cells expressing the hDAT-PG^584,585^AA after preincubation with ibogaine analogs. ***A***. Comparison of [^3^H]DA uptake in HEK293 cells transiently expressing wild type (WT) hDAT or hDAT-PG^584,585^AA. Each symbols represents the result from an individual experiment (done in triplicate); means± S.D. are also shown. ***B***. hDAT-PG^584,585^AA expressing cells were incubated in the presence of the indicated compounds (30 μM in all instances but bupropion, 100 μM). After 24 h, specific [^3^H]dopamine uptake was determined as outlined under “Methods”. Values from individual experiments (done in triplicate) are shown as dots; box plots show the median and the interquartile range. ***C***. *(Top)* Concentration-response curves for pharmacochaperoning by the indicated compounds (selected as positive hits from *panel B).* Rescued uptake was normalized to that achieved by 30 μM noribogaine (=100%) to account for inter-experimental variations. Rescue by **9b** was plotted separately because of its very high efficacy in comparison to other analogs *(bottom).* The solid lines were drawn by fitting the data to the equation for a rectangular hyperbola (for EC_50_ and E_max_ of mutant rescue see **Table 1**). Data were obtained in at least three independent experiments carried out in triplicate. The error bars indicate S.D.

### Rescue of DAT-PG AA in flies by *in vivo* pharmacochaperoning with the ibogaine analog 9b

In drosophila, dopaminergic projections into the fan-shaped body are required to maintain a wake/sleep cycle *(Liu et al., 2012; Pimentel et al., 2016).* In the absence of functional DAT, flies are hyperactive and have abnormal sleep regulation. Accordingly, drosophilae, in which the endogenous DAT gene is disrupted, are referred to as *fumin* (i.e., Japanese for sleepless) flies *(Kume et al., 2005).* We previously verified that noribogaine was an effective pharmacochaperone *in vivo* for some folding-deficient versions of DAT in flies: when expressed in dopaminergic neurons of *fumin* flies, folding-deficient mutants of human DAT were retained in the ER of the soma of PPL1 neurons but delivered to the axonal terminals of the fan-shaped body, if flies were administered noribogaine; concomitantly, sleep was restored *(Kasture et al., 2016; Asjad et al., 2017).* Accordingly, we tested compound **9b** to determine if it was also effective as a pharmacochaperone *in vivo*. We generated *fumin* flies, which harbored the human cDNA encoding DAT-PG^584,585^AA under the control of GAL4 driven from a tyrosine hydroxylase promoter *(Friggi-Grelin et al., 2003).* Adult flies were individually placed in transparent tubes, which contained the food pellet supplemented with designated concentrations of compound **9b** or of noribogaine, and allowed to recover for one day; they then spent an additional day of a 12h light/12h dark cycle (marked as yellow and black rectangle in **Fig. 4A**) to entrain their circadian rhythm. It is evident that, when subsequently released into a dark-dark cycle (marked as gray and black rectangle in **Fig. 4A**), these flies were as hyperactive as *fumin* flies (*cf.* green and red symbols in **Fig. 4A**). Their hyperactivity was greatly reduced by treatment of either compound **9b** or noribogaine at a concentration of 30 and 100 μM, respectively, in the food pellet *(cf.* blue and orange symbols in **Fig. 4A**). In fact, both compound **9b** and noribogaine reduced the locomotor activity to that seen in the isogenic control line w1118. We quantified the effect of increasing doses of compound **9b** (administered by raising its concentration in the food pellet from 10 to 100 μM) on sleep time (**Fig. 4B**): on average this increased in a dose-dependent manner by about 2.5-fold, i.e. from 300 min in the absence of any pharmacochaperone in the food pellet to 750 min with 100 μM of compound **9b** in the food pellet. Sleep duration was comparable to that seen after administration of feed containing 100 μM noribogaine and approached that seen in w1118 flies. This indicates that pharmacochaperoning by compound **9b** rescued DAT-PG^584,585^AA in amounts sufficient to restore clearance of dopamine from the synapse and to thus reinstate normal dopaminergic transmission. Finally, the effect of compound **9b** (and of noribogaine) was specific: sleep duration was neither affected by compound **9b** (or noribogaine) in w1118 flies harboring an intact DAT nor in DAT-deficient fumin flies (**Fig. 4C**).

**Figure 4.**
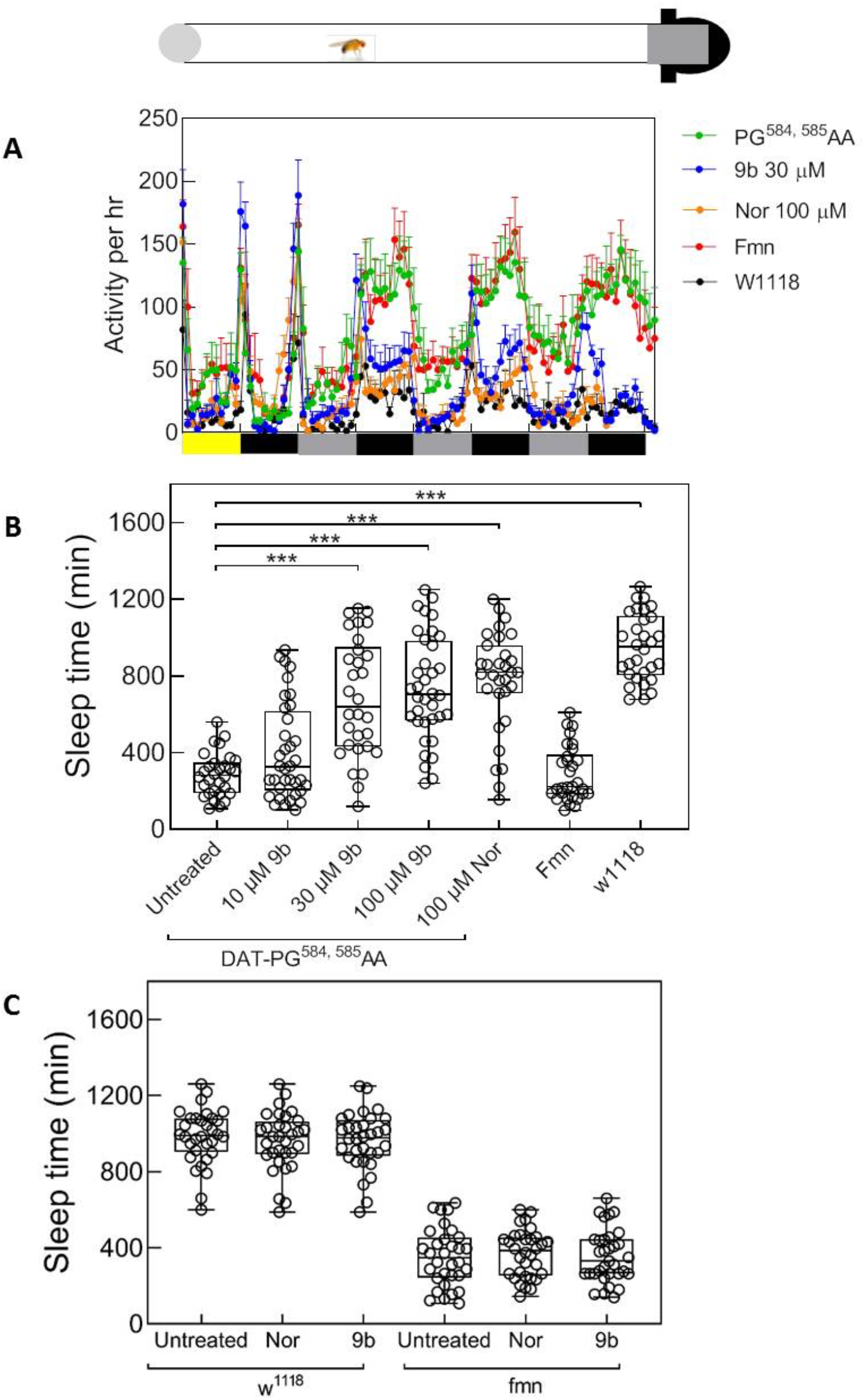
Pharmacochaperones restore sleep in flies harboring DAT-PG^584,585^AA. ***A.*** Locomotor activity (measured over 1 min interval and binned into 60 min intervals) was recorded from day 2 to day 5 using a Drosophila activity monitor system (schematically represented above the graph, for details see Methods). The data are means ± SEM from three independent experiments, which were each carried out in parallel with at least 10 flies/condition. The diurnal light cycles (light-dark on day 2, and dark-dark on days 3-5) are outlined schematically below the graph. ***B.*** and **C,** sleep time of treated DAT-PG^584,585^AA mutant flies. Flies expressing hDAT-PG^584,585^AA in the *fumin* background were fed with food pellets containing the indicated concentrations of noribogaine and 9b. ***C.*** w1118 and fumin flies were used as control and were given food containing 100 μM noribogaine or 100 μM 9b. Locomotor activity on day 4 was used to quantify the sleep time using pySolo software. Empty circle represents individual flies. Statistical significance of the observed differences was determined by analysis of variance followed by Dunn’s post-hoc test (***, p < 0.001).

### Pharmacochaperoning of folding-deficient DAT-mutants associated with human disease by the ibogaine analog 9b

The efficacy of compound **9b**, which was seen *in vivo*, is promising. We therefore examined the ability of compound **9b** to rescue all disease-relevant mutants of DAT. When heterologously expressed in HEK293 cells, measurable substrate influx was restored in six mutants (**Fig. 5**); in four of these mutants, i.e. DAT-G^386^R, -R^521^W, -P^395^L, and -P^554^L, compound **9b** was more efficacious than the reference compounds ibogaine and noribogaine. In contrast, preincubation in the presence of the corresponding deconstructed analogs **3c** and **4c** invariably failed to restore any transport activity. Compound **9b** did not rescue the function of DAT-R^85^L, -V^158^F, -L^224^P, -G^327^R, -L^368^Q, -Y^470^S and -P^529^L in any appreciable manner (data not shown).

**Figure 5.**
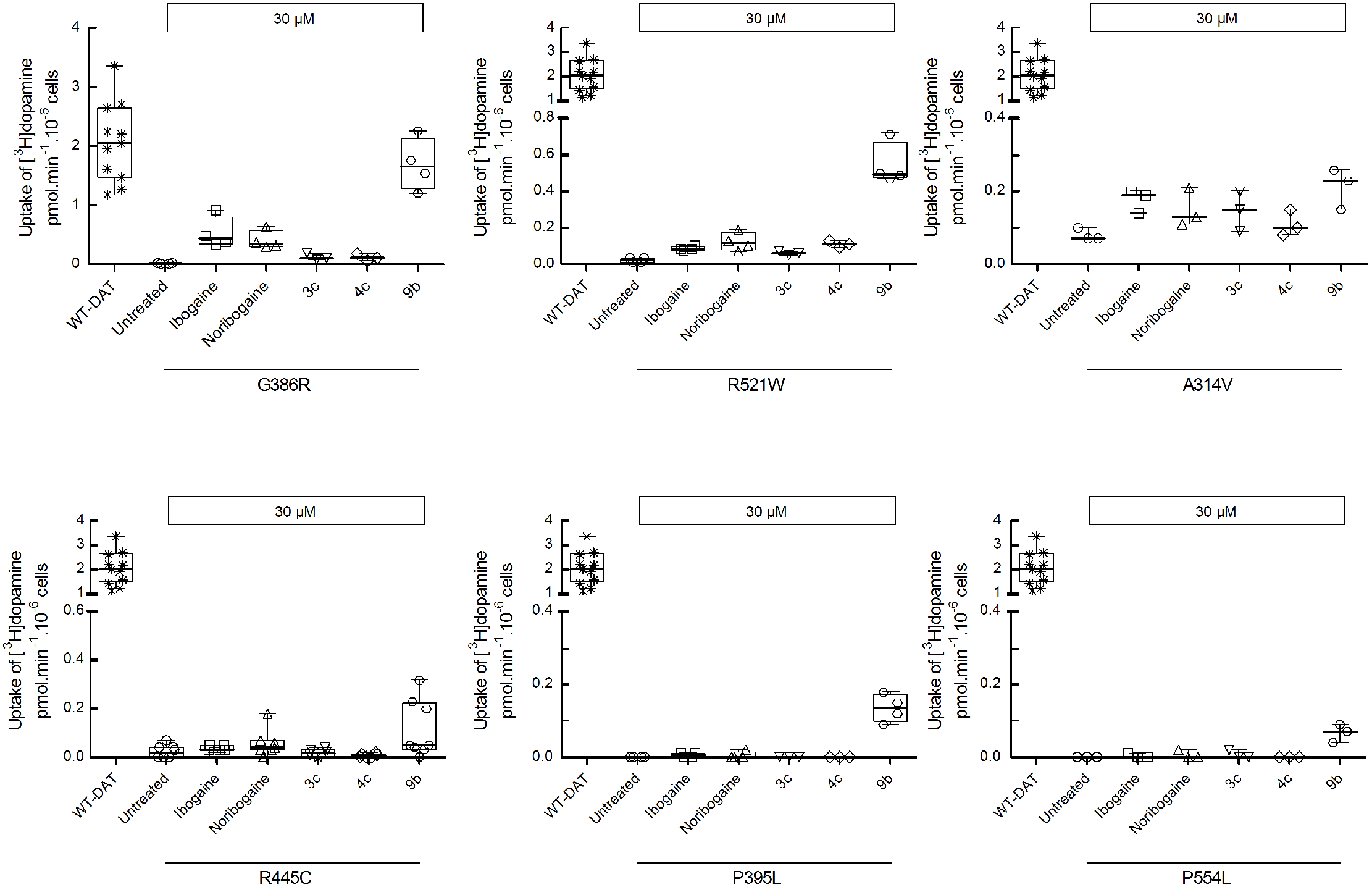
Pharmacochaperoning of DTDS-associated DAT mutants by 9b. HEK293 cells were transiently transfected with plasmids encoding YFP-tagged wild-type hDAT or each of the 13 hDAT mutants discovered in patients suffering from dopamine transporter deficiency syndrome (DTDS). After 24 h, the cells were seeded onto 96-well plates for 24 h and were either incubated in the absence (untreated) or presence of either of ibogaine, noribogaine or of the fluorinated ibogaine analogs **3c**, **4c** and **9b** (all at 30 μM) for another 24 h. The cells were washed 4 times with Krebs-MES buffer (pH 5.5) and once with Krebs-HEPES buffer (pH 7.4) to completely remove extracellular reservoir of the compounds. Uptake of [^3^H]dopamine was subsequently measured as outlined under “Material and Methods.” The mutants with positive pharmacochaperoning effect by **9b** are represented in this figure. The data are represented as the individual values from at least three independent experiments carried out in triplicate for wild-type hDAT and the DAT mutants, and as box plots with the median and the interquartile range; whiskers indicate the 95% confidence interval. WT-DAT uptake values were used as a control for transfection efficiency. Compound **9b** failed to restore activity in the other DAT mutants (i.e. DAT-R^85^L, -V^158^F, -L^224^P, -G^327^R, -L^368^Q, -Y^470^S, and -P^529^L).

## Discussion

More than sixty point mutations have been identified, which give rise to human disease, because they lead to misfolding of a member of the SLC6 transporter family *(Freissmuth et al., 2018).* Folding defects can be overcome by pharmacochaperones, which reduce energy barriers in the folding trajectory: stalled intermediates proceed to reach the native fold. Pharmacochaperoning by ibogaine was serendipitously discovered, when studying ER export of SERT *(El-Kasaby et al., 2010).* Ibogaine and its metabolite noribogaine effectively rescue some disease-relevant DAT mutants, but with limited efficacy *(Beerepoot et al., 2016; Asjad et al., 2017).* Progress is contingent on understanding the attributes, which account for the pharmacochaperoning action of ibogaine. Here, we relied on an orthogonal approach by interrogating the chemical space populated by variations of the ibogaine structure and by probing their effect on the folding space of SERT and DAT, using two analogous mutations and the disease-associated mutations of DAT. Our observations lead to the insight that affinity for the wild type transporter (i.e. the native folded state) and pharmacochaperoning activity are not tightly linked, as the structure-activity relationships differ. This conclusion is based on the following lines of evidence: the affinity for SERT is governed predominantly by substitution on the indole ring with structural rigidity playing a minor role (**Table 1**). Accordingly, hydroxy-(**3f**, **9d**) and fluoro-(**3c**, **4c**, **9b**) analogs had higher affinities for SERT than methoxy- and unsubstituted analogs, but **9d** had lower affinity for SERT than its more flexible analogue **3f**.

The substitution on the indole ring is less important a determinant for DAT affinity. The most important determinant for pharmacochaperoning efficacy is the structural rigidity imparted by the azepine ring, which is retained by ibogaine, noribogaine and compounds of the **9a**–**9d** series. In contrast, this rigidity is not required for supporting moderate to high-affinity binding to SERT and DAT. The fluorinated compounds **3c** and **4c**, for instance, were as potent as **9b** in inhibiting SERT, but they were substantially less efficacious and less potent in rescuing SERT-PG^601,602^AA. Similarly, these three analogs only differed modestly in their affinity for DAT, but only compound **9b** was an efficacious pharmacochaperone for DAT-PG^584,585^AA. In fact, compound **9b** meets three important criteria to be considered a significant advance in the pharmacochaperoning of DAT mutants: (i) the pharmacochaperoning efficacy of compound **9b** exceeded that of the parent compound noribogaine. (ii) Compound **9b** had a therapeutic window *in vivo,* i.e. there was an effective dose range, where it restored sleep in drosophilae harboring DAT-PG^584,585^AA. This effect was specifically linked to the pharmacochaperoning action, because compound **9b** did not affect locomotion and sleep in control flies (harboring endogenous DAT) or *fumin* flies lacking DAT. (iii) Compound **9b** restored folding to disease-associated DAT mutants, which were unresponsive to noribogaine, thus doubling the number of DAT mutants potentially amenable to folding correction.

The mechanistic basis underlying pharmacochaperoning is poorly understood. At least four mechanisms are conceivable, i.e. (i) stabilization of the native state or (ii) of a folding intermediate, (iii) prevention of aggregate formation or (iv) dissolution of aggregates *(Marinko et al., 2019).* Stabilization of the native state is posited to be the most common mechanism of action for pharmacochaperones. In this instance, chaperoning efficacy is related to binding affinity *(Marinko et al., 2019).* Our observations are difficult to reconcile with binding to the native state, because (i) the structure-activity relationships for binding to the wild type transporters – i.e. the native state – differed substantially from for pharmacochaperoning. (ii) The EC_50_-values for rescuing DAT-PG^584,585^AA and SERT-PG^601,602^AA differed by 400-fold. This difference was substantially larger than variation in affinity for the native state of SERT and DAT. This indicates that compound **9b** has a high and low affinity for the relevant folding intermediate(s) of SERT-PG^601,602^AA and of DAT-PG^584,585^AA, respectively. (iii) The relative pharmacochaperoning efficacy of ibogaine, noribogaine and compound **9b** was highly dependent on the nature of the DAT mutant: in DAT-PG^584,585^AA and in several disease-relevant DAT mutants (i.e., DAT-G^386^R, -R^521^W, – P^395^L, P^554^L) compound **9b** was substantially more efficacious than ibogaine and noribogaine, but this was not the case in DAT-A^314^V and DAT-R^445^C. This variation in relative efficacy is again difficult to reconcile with stabilization of the native state. (iv) Circumstantial evidence also argues that the other two proposed mechanisms of pharmacochaperoning – inhibition of aggregation and disassembly of aggregates – do not apply: diseases arising from mutations in SLC6 transporter can be transmitted in both a recessive and a dominant fashion. Because SLC6 transporters are exported from the ER in an oligomeric form, folding-deficient mutants can act in a dominant-negative manner and retain the wild type transporter (*Chiba et al., 2014; Lopez-Corcuera et al., 2019).* However, all disease-relevant human DAT mutants are transmitted as recessive alleles *(Kurian et al., 2011; Ng et al., 2014).* Thus, their folding trajectory is stalled at a stage, where they are complexed with and shielded by ER chaperones such as calnexin, which precludes oligomerization *(Korkhov et al., 2008).* Taken together, these observations indicate that binding to folding intermediate(s) is the most likely mechanism underlying the pharmacochaperoning action of compound **9b**.

Individual folding-deficient mutants of SERT and DAT are stalled at different points of their folding trajectory and differ in their susceptibility to pharmacochaperoning *(El-Kasaby et al., 2014; Koban et al., 2015; Beerepoot et al., 2016; Asjad et al., 2017; Freissmuth et al., 2017).* Our approach relied on using the two analogous folding deficient mutants SERT-PG^601,602^AA and DAT-PG^584,585^AA by assuming that these two proteins are stalled at related positions of their folding trajectory. However, compound **9b** was more than 100-fold more potent in rescuing SERT-PG^601,602^AA than DAT-PG^584,585^AA. A similar discrepancy was seen in the naphthylamine series of pharmacochaperones *(Bhat et al., 2017):* PAL-287 was more effective than PAL-1045 in rescuing DAT-PG^584,585^AA, the reverse was true for SERT-PG^601,602^AA (cf. **Fig. 3** and *Bhat et al., 2017*). Taken together, these observations are again consistent with the conjecture that the compounds act as pharmacochaperones by binding to folding intermediates rather than by stabilizing the native state. The hypothetical model posits that, in the folding trajectory of wild type SLC6 transporters, there is a large isoenergetic conformational search space. Mutations convert this smooth surface into a rugged landscape. Accordingly, they create multiple traps, which reside at different locations in the peripheral ring of the champagne glass-like energy landscape *(Dill and Chan, 1997).* Some of these traps can be overcome by pharmacochaperoning, but they differ – even for closely related folding-deficient mutations – in their position in the conformational search space (*El-Kasaby et al., 2014*). Hence, the folding intermediates in the vicinity of the trap(s) differ in their ability to bind to and respond to a pharmacochaperoning ligand. General rules, which govern folding of helical membrane proteins, have been inferred but the details are obscure *(Chiba et al., 2014; Marinko et al., 2019).* Progress is hampered, because the nature of the folding intermediate(s) is poorly understood. The affinity of compound **9b** for the folding intermediate(s) of SERT-PG^601,602^AA was estimated to be in the submicromolar range. Hence, compound **9b** may be useful as a starting point to develop probes to address the folding trajectory of SERT-PG^601,602^AA and other misfolded SERT variants. We anticipate that the resulting insights will also advance the search for additional pharmacochaperones. These are needed, because the majority of disease-associated DAT mutants are still not remedied by the available pharmacochaperones. Moreover, compound **9b** is a significant breakthrough, because it not only expands the number of rescued DAT mutants but it also restores the functional activity of DAT-G386R essentially to wild type levels. Thus, compound **9b** provides a proof-of-principle that it is possible to fully correct the folding defect of a mutant by pharmacochaperoning.

## Material and Methods

### Cell culture and materials

Cells were propagated Dulbecco’s Modified Eagle Medium (DMEM) supplemented with 10% heat-inactivated fetal bovine serum (FBS), 100 u · 100mL^-1^ penicillin and 100 u · 100mL^-1^ streptomycin. Medium used in the maintenance of stable lines (see below) was, in addition, supplemented with 50 μg ml^-1^ geneticin (G418) for selection. HEK293 cells were transfected by combining plasmid DNA with PEI (linear 25 kDa polyethylenimine; Santa Cruz, SC-360988A) at a ratio of 1:3 (w:w) in serum-free DMEM. The plasmids encoding YFP-tagged human wild type and mutant versions of SERT (El-Kasaby et al., 2014) and of DAT (Kasture et al.,2016; Asjad et al., 2017) were previously described; YFP-tagged DAT-PG^584,585^AA was created by introducing the substitutions with created with the QuikChange Lightning Site-Directed Mutagenesis Kit (Stratagene, La Jolla, CA), using wild type YFP-fagged DAT as the template. The mutations were confirmed by automatic DNA sequencing (LGC Labor GmbH Augsburg, Germany). For the pharmacochaperoning experiments, noribogaine was purchased from Cfm Oskar Tropitzsch GmbH (Marktredwitz, Germany). [^3^H]5-HT (serotonin, 41.3 Ci/mmol), and [^3^H]dopamine (DA, 39.6 Ci/mmol) were purchased from PerkinElmer Life Sciences. Scintillation mixture (Rotiszint^®^ eco plus) was purchased from Carl Roth GmbH (Karlsruhe, Germany). Cell culture media and antibiotics were obtained from Sigma and Invitrogen, respectively. Anti-GFP antibody (rabbit, ab290) was from Abcam (Cambridge, UK). An antibody raised against an N-terminal peptide of the G protein β subunit was used to verify comparable loading of lanes *(Hohenegger et al., 1996).* Horseradish peroxidase–linked anti-rabbit IgG1 antibody was purchased from Amersham Biosciences. All other chemicals were of analytical grade.

### Radioligand Binding Studies

For DAT binding assays, frozen striata, previously dissected from freshly harvested male Sprague-Dawley rat brains (supplied on ice by Bioreclamation, Hicksville, NY), were homogenized in 20 volumes (w/v) of ice-cold modified sucrose phosphate buffer (0.32 M sucrose, 7.74 mM Na_2_HPO_4_, and 2.26 mM NaH_2_PO_4_, pH adjusted to 7.4) using a Brinkman Polytron (Setting 6 for 20 s) and centrifuged at 48,400 x g for 10 min at 4°C. The resulting pellet was washed by resuspension in buffer, the suspension was centrifuged again, and the final pellet resuspended in ice-cold buffer to a concentration of 20 mg/mL (original wet weight/volume, OWW/V). Experiments were conducted in 96-well polypropylene plates containing 50 μL of various concentrations of the inhibitor, diluted using 30% DMSO vehicle, 300 μL of sucrose phosphate buffer, 50 μL of [^3^H]WIN 35,428 (final concentration 1.5 nM; *KD* = 28.2 nM; PerkinElmer Life Sciences, Waltham, MA), and 100 μL of tissue (2.0 mg/well OWW). All compound dilutions were tested in triplicate and the competition reactions started with the addition of tissue, and the plates were incubated for 120 min at 0-4°C. Nonspecific binding was determined using 10 μM indatraline.

For SERT binding assays, frozen brain stem tissue, previously dissected from freshly harvested male Sprague-Dawley rat brains (supplied on ice by Bioreclamation, Hicksville, NY), was homogenized in 20 volumes (w/v) of 50 mM Tris buffer (120 mM NaCl and 5 mM KCl, adjusted to pH 7.4) at 25°C using a Brinkman Polytron (at setting 6 for 20 s) and centrifuged at 48,400 x g for 10 min at 4°C. The resulting pellet was resuspended in buffer, the suspension was centrifuged, and the final pellet suspended in buffer again to a concentration of 20 mg/mL (OWW/V). Experiments were conducted in 96-well polypropylene plates containing 50 μL of various concentrations of the inhibitor, diluted using 30% DMSO vehicle, 300 μL of Tris buffer, 50 μL of [^3^H]citalopram (final concentration 1.5 nM; *K_d_* = 6.91 nM; PerkinElmer Life Sciences, Waltham, MA), and 100 μL of tissue (2.0 mg/well OWW). All compound dilutions were tested in triplicate and the competition reactions started with the addition of tissue, and the plates were incubated for 60 min at rt. Nonspecific binding was determined using 10 μM fluoxetine.

For all binding assays, incubations were terminated by rapid filtration through Perkin Elmer Uni-Filter-96 GF/B presoaked in either 0.3% (SERT) or 0.05% (DAT) polyethylenimine, using a Brandel 96-well harvester manifold or Brandel R48 filtering manifold (Brandel Instruments, Gaithersburg, MD). The filters were washed a total of 3 times with 3 mL (3 × 1 mL/well or 3 x 1 mL/tube) of ice cold binding buffer. Perkin Elmer MicroScint 20 Scintillation Cocktail (65 μL) was added to each filter well. Radioactivity was counted in a Perkin Elmer MicroBeta Microplate Counter. IC_50_ values for each compound were determined from inhibition curves and K_i_ values were calculated using the Cheng-Prusoff equation. When a complete inhibition could not be achieved at the highest tested concentrations, *K*_i_ values were estimated by extrapolation after constraining the bottom of the dose-response curves (= 0% residual specific binding) in the non-linear regression analysis. These analyses were performed using GraphPad Prism version 8.00 for Macintosh (GraphPad Software, San Diego, CA). K_d_ values for the radioligands were determined via separate homologous competitive binding or radioligand binding saturation experiments. *K_i_* values were determined from at least 3 independent experiments performed in triplicate and are reported as means ± S.D.

### [^3^H]5-HT and [^3^H]dopamine uptake assays

For uptake inhibition assays, HEK293 cells stably expressing either wild-type human YFP-tagged hSERT or YFP-tagged hDAT were seeded on poly-D-lysine-coated 96-well plates at a density of ~20,000 cells/well. After 24 h, the medium in each well was aspirated and the cells were washed once with Krebs-HEPES buffer (10 mM HEPES.NaOH, pH 7.4, 120 mM NaCl, 3 mM KCl, 2 mM CaCl_2_, 2 mM MgCl_2_, and 2 mM glucose). Cells were pre-incubated in buffer containing logarithmically spaced concentrations (0.003–300 μM) of ibogaine analogs for 10 minutes. Subsequently the reaction was started by by addition of substrate (0.4 μM of either [^3^H]5-HT or [^3^H]dopamine) at constant concentrations of ibogaine analogs for 1 minute. The reaction was terminated by aspirating the reaction medium followed by a wash with ice-cold buffer. The cells were lysed with 1% SDS and the released radioactivity was quantified by liquid scintillation counting.

For uptake assays determining functional rescue of mutant transporters, HEK293 cells were transfected with either YFP-tagged SERT-PG^601,602^AA or YFP-tagged DAT-PG^584,585^AA plasmids. Transfected cells were seeded on poly-D-lysine-coated 96-well plates at a density of ~60-80,000 cells/well either in the absence or presence of increasing concentrations (0.1 – 100 μM) of the ibogaine analogs. After 24 h, the cells were washed four times with Krebs-MES buffer (10 mM 2-(N-morpholino)ethanesulfonic acid, pH 5.5, 120 mM NaCl, 3 mM KCl, 2 mM CaCl_2_, 2 mM MgCl_2_, and 2 mM glucose) in a 10 min interval and once with Krebs-HEPES (pH 7.4) buffer. The cells were subsequently incubated with 0.2 μM of [^3^H]5-HT for 1 minute or [^3^H]dopamine for 5 min. and processed as outlined above.

### Immunoblotting after pharmacochaperoning of SERT-PG^601,602^AA or DAT-PG^584,585^AA

HEK293 cells were transiently transfected with plasmids encoding either wild type SERT, SERT-PG^601,602^AA, wild type DAT or DAT-PG^584,585^AA. Approximately 1.5 – 2 x 10^6^ of these transfected cells were seeded either in 6-well plates or 6 cm dishes in the presence of 30 μM of the individual candidate hits identified by uptake assays. After 24 h, cells were washed thrice with ice-cold phosphate-buffered saline, detached by mechanical scraping, and harvested by centrifugation at 1000 x g for 5 min. The cell pellet was lysed in a buffer containing Tris·HCl, pH 8.0, 150 mm NaCl, 1% dodecyl maltoside, 1 mm EDTA, and protease inhibitors (Complete™, Roche Applied Science). This soluble protein lysate was separated from the detergent-insoluble material by centrifugation (16,000 × g for 15 min at 4 °C). An aliquot of this lysate (20 μg) was mixed with 1% SDS and 20 mM DTT containing sample buffer, denatured at 45 °C for 30 min, and resolved in denaturing polyacrylamide gels. After protein transfer onto nitrocellulose membranes, the blots were probed with an antibody against GFP (rabbit, ab290) at a 1:3000 dilution overnight. This immunoreactivity was detected using a horseradish peroxidase conjugated secondary antibody (1:5000, Amersham ECL Prime Western Blotting Detection Reagent). In separate experiments, lysates were prepared from cells treated in the absence or presence of 30 μM DG4-69; these were incubated in the presence and absence of endoglycosidase H (New England Biolabs) (16 = Asjad et al., 2017) and aliquots (20 μg) were then resolved electrophoretically as described above. Densitometric analyses of individual blots were done using ImageJ.

### Fly genetics, treatment and locomotion assay

The transgenic UAS reporter line for YFP-tagged hDAT-PG^584, 585^AA was generated using pUASg-attB vector (gift from Drs. Bischof and Basler, University of Zürich). The sequenced construct was injected into embryos from ZH-86Fb flies (Bloomington stock no 24749). Positive transformants were selected and crossed with balancer flies (Bloomington stock no. 3704). Fumin flies (fmn or DAT-KO mutant flies) was a generous gift from Dr. Kume, Nagoya City University, Japan. Tyrosine hydroxylase Gal4 (TH-Gal4, Bloomington stock no. 8848) was used to drive the expression of hDAT-PG^584, 585^AA in dopaminergic neurons. Isogenized *fmn* and *w^1118^* flies were used as control. The genotypes of flies used in Fig. 7a and b were *w^1118^; fmn(w; roo{}DAT^fmn^); TH-Gal4/UAS-hDAT-PG^584, 585^AA* (PG^584, 585^AA)*, w^1118^/y; fmn(w; roo{}DAT^fmn^); +/+* (Fmn) and *w^1118^.* All flies were kept at 25 °C in a 12-h light/12-h dark cycle, and all crosses were performed at 25 °C. As described previously (14 = Kasture et al., 2016), locomotion assay was performed on three-to-five-day old male flies using *Drosophila* activity monitor system (DAM2, Trikinetics, Waltham, MA). Briefly, individual flies were housed in 5-mm-diameter glass tubes carrying food pellet supplemented with specified concentrations of noribogaine and **9b**. Flies were entrained for 12h:12h day: night rhythm for first two days and locomotion activity was studied on second day in subsequent 12h; 12h dark: dark phase. Locomotion activity was measured in 1 min bins and pySolo software (Gilestro and Cirelli, 2009) was used to quantify sleep time.

## Supporting Information

### Synthesis

All chemicals and solvents were purchased from chemical suppliers unless otherwise stated and used without further purification. ^1^H and ^13^C NMR spectra were acquired using a Varian Mercury Plus 400 spectrometer at 400 MHz and 100 MHz, respectively. Chemical shifts are reported in parts-per-million (ppm) and referenced according to deuterated solvent for ^1^H NMR spectra (CDCl_3_, 7.26; D_2_O, 4.79 or DMSO-d_6_, 2.50) and ^13^C NMR spectra (CDCl_3_, 77.2 or DMSO-d_6_, 39.52). Gas chromatography-mass spectrometry (GC/MS) data were acquired (where obtainable) using an Agilent Technologies (Santa Clara, CA) 7890B GC equipped with an HP-5MS column (cross-linked 5% PH ME siloxane, 30 m × 0.25 mm i.d. × 0.25 μm film thickness) and a 5977B mass-selective ion detector in electron-impact mode. Ultrapure grade helium was used as the carrier gas at a flow rate of 1.2 mL/min. The injection port and transfer line temperatures were 250 and 280 °C, respectively, and the oven temperature gradient used was as follows: the initial temperature (70 °C) was held for 1 min and then increased to 300 °C at 20 °C/min over 11.5 min, and finally maintained at 300 °C for 4 min. All column chromatography was performed using a Teledyne Isco CombiFlash RF flash chromatography system. Combustion analyses were performed by Atlantic Microlab, Inc. (Norcross, GA) and agree with ± 0.4% of calculated values. HRMS (mass error within 5 ppm) and MS/MS fragmentation analysis were performed on a LTQ-Orbitrap Velos (Thermo-Scientific, San Jose, CA) coupled with an ESI source in positive ion mode to confirm the assigned structures and regiochemistry. All melting points were determined on an OptiMelt automated melting point system and are uncorrected. On the basis of NMR and combustion data, all final compounds are >95% pure.

*3-(2-(8-azabicyclo[3.2.1]octan-8-yl)ethyl)-1H-indole (**3a**).* 8-Azabicyclo[3.2.1]octane (**1**) (222.4 mg, 2 mmol), 3-(2-bromoethyl)-1H-indole (448.2 mg, 2 mmol), K_2_CO_3_ (1.1 g, 8 mmol) and acetonitrile (24 mL) were added in a sealed bottle (100 mL). The reaction mixture was stirred at 100 °C overnight and filtered. The filtrate was evaporated and purified by flash column chromatography (DCM/MeOH/NH4OH = 95: 5: 0.5) to give the product (470 mg, 92% yield) as a yellow oil. The free base was converted to the HCl salt and recrystallized from methanol to give a white solid. Mp 247-248 °C; GC/MS (EI) *m/z* 254 (M^+^); ^1^H NMR (400 MHz, CDCl_3_) δ 8.05 (s, 1H), 7.62-7.63 (m, 1H), 7.34-7.36 (m, 1H), 7.02-7.19 (m, 3H), 3.34 (m, 2H), 2.95-2.99 (m, 2H), 2.68-2.72 (m, 2H), 1.94-1.99 (m, 2H), 1.76-1.85 (m, 2H), 1.35-1.64 (m, 6H); ^13^C NMR (100 MHz, CDCl_3_) δ 136.2, 127.6, 121.9, 121.4, 119.2, 118.9, 114.9, 111.1, 59.6, 53.4, 30.7, 26.5, 25.0, 16.7,; Anal. (C_17_H_22_N_2_ · HCl) C, H, N.

*3-(3-(8-azabicyclo[3.2.1]octan-8-yl)propyl)-1H-indole (**3b**).* 3-(1H-indol-3-yl)propanoic acid (378.4 mg, 2 mmol) was dissolved in THF (20 mL), CDI (1 equiv) was added and stirred for 2 h at rt followed by adding 8-azabicyclo[3.2.1]octane (**1**) (222.4 mg, 2 mmol) in THF (13 mL). The reaction mixture was stirred overnight at rt. The solvent was removed *in vacuo* and residue was diluted with CHCl_3_ (50 mL) and washed with saturated aq Na_2_CO_3_ solution (2 x 30 mL). The organic layer was dried with MgSO_4_ and concentrated *in vacuo.* The crude product was purified by column chromatography (DCM/MeOH/NH_4_OH = 97: 3: 0.5) to give the amide as yellow oil. The amide was dissolved in anhydrous THF (5.6 mL) and added dropwise to the suspension of LAH (110 mg, 2.84 mmol) in THF (2 mL) at 0 °C. The reaction mixture was allowed to warm to rt and stirred for 3 h. H_2_O (0.3 mL) was added carefully at 0 °C, followed by the addition of 0.5 mL of aq NaOH (2 M). The resulting mixture was filtered, and the filtrate was dried (K_2_CO_3_) filtered and the solvent was removed *in vacuo*. The residue was purified by column chromatography (DCM/MeOH/NH4OH = 95: 5: 0.5) to give the product (250 mg, 47% yield over two steps) as a yellow oil. The free base was converted to the HCl salt and recrystallized from methanol to give a tan foam; GC/MS (EI) *m/z* 268 (M^+^); ^1^H NMR (400 MHz, CDCl_3_) δ 7.98 (s, 1H), 7.60-7.62 (m, 1H), 7.33-7.35 (m, 1H), 6.99-7.19 (m, 3H), 3.20 (m, 2H), 2.76-2.80 (m, 2H), 2.42-2.46 (m, 2H), 1.87-1.95 (m, 4H), 1.41-1.78 (m, 6H), 1.26-1.33 (m, 2H); ^13^C NMR (100 MHz, CDCl_3_) δ 136.3, 127.6, 121.8, 121.0, 119.0, 116.7, 111.0, 59.4, 52.3, 30.7, 29.2, 26.4, 23.1, 16.8; Anal. (C_18_H_24_N_2_ · HCl · 0.5H_2_O · 0.25 *i*-PrOH) C, H, N.

*3-(2-(8-azabicyclo[3.2.1]octan-8-yl)ethyl)-5-fluoro-1H-indole (**3c**).* Compound **3c** was prepared as described for **3b** using 8-azabicyclo[3.2.1]octane (**1**) (111.2 mg, 1 mmol) and 3-(5-fluoro-1H-indol-3-yl)propanoic acid (207.2mg, 1 mmol) to give the product (250 mg, 28% yield over two steps) as a yellow oil. GC/MS (EI) *m/z* 286 (M^+^); ^1^H NMR (400 MHz, CDCl_3_) δ 8.58 (s, 1H), 7.21-7.24 (m, 2H), 6.89-6.96 (m, 2H), 3.22-3.24 (m, 2H), 2.70-2.74 (m, 2H) 2.43-2.47 (m, 2H), 1.73-1.94 (m, 6H), 1.30-1.58 (m, 6H); ^13^C NMR (100 MHz, CDCl_3_) δ 158.7, 156.4, 132.8, 128.1, 128.0, 123.0, 116.7, 116.6, 111.6, 111.5, 110.1, 110.0, 103.9, 103.6, 59.5, 52.2, 30.6, 29.1, 26.4, 23.0, 16.7; Anal. (C_18_H_23_FN_2_ · 0.5H_2_O) C, H, N.

*3-(2-(8-azabicyclo[3.2.1]octan-8-yl)ethyl)-5-methoxy-1H-indole (**3d**).* Compound **3d** was prepared as described for **3b** using 8-azabicyclo[3.2.1]octane (**1**) (222.4 mg, 2 mmol) and 2-(5-methoxy-1H-indol-3-yl)acetic acid (410.0 mg, 2 mmol) to give the product (380 mg, 49% yield over two steps) as a brown oil. The free base was converted to the HCl salt and recrystallized from methanol to give a tan foam; GC/MS (EI) *m/z* 284 (M^+^); ^1^H NMR (400 MHz, CDCl_3_) δ 7.89 (s, 1H), 7.22-7.24 (m, 1H), 7.07 (s, 1H), 7.00 (s, 1H), 6.83-6.86 (m, 1H), 3.86 (s, 3H), 3.34-3.35 (m, 2H), 2.91-2.95 (m, 2H), 2.67-2.70 (m, 2H), 1.95-1.99 (m, 2H), 1.78-1.85 (m, 2H), 1.46-1.61 (m, 6H); ^13^C NMR (100 MHz, CDCl_3_) δ 153.9, 131.3, 128.0, 122.2, 114.6, 112.1, 111.8, 100.8, 59.6, 55.9, 53.3, 30.7, 26.5, 25.0, 16.7; Anal. (C_18_H_24_N_2_O · HCl · H_2_O) C, H, N.

*3-(3-(−8-azabicyclo[3.2.1]octan-8-yl)propyl)-5-methoxy-1H-indole (**3e**).* Compound **3e** was prepared as described in for **3b** using 8-azabicyclo[3.2.1]octane (**1**) (222.4 mg, 2 mmol) and 3-(5-methoxy-1H-indol-3-yl)propanoic acid (438.0 mg, 2 mmol) to give the product (380 mg, 79% yield over two steps) as a tan oil; GC/MS (EI) *m/z* 298 (M^+^); ^1^H NMR (400 MHz, CDCl_3_) δ 8.07 (s, 1H), 7.21-7.24 (m, 1H), 7.04 (s, 1H), 6.95 (s, 1H), 6.83-6.94 (m, 1H), 3.86 (s, 3H), 3.21-3.23 (m, 2H), 2.73-2.76 (m, 2H), 2.44-2.48 (m, 2H), 1.87-1.94 (m, 4H), 1.72-1.80 (m, 2H), 1.43-1.59 (m, 4H), 1.30-1.34 (m, 2H); ^13^C NMR (100 MHz, CDCl_3_) δ 153.8, 131.5, 128.0, 121.9, 116.4, 112.0, 111.7, 100.9, 59.4, 56.0, 52.3, 30.7, 29.1, 26.4, 23.1, 16.7; Anal. (C_19_H_26_N_2_O · 0.5H_2_O) C, H, N.

*3-(2-(8-azabicyclo[3.2.1]octan-8-yl)ethyl)-1H-indol-5-ol (**3f**).* Compound **3f** was prepared as described for **3b** using 8-azabicyclo[3.2.1]octane (**1**) (222.4 mg, 2 mmol) and 2-(5-hydroxy-1H-indol-3-yl)acetic acid (382.4 mg, 2 mmol) to give the product (158 mg, 29% yield over two steps) as a tan oil; GC/MS (EI) *m/z* 270 (M^+^); ^1^H NMR (400 MHz, CDCl_3_) δ 8.12 (s, 1H), 7.11-7.13 (m, 1H), 6.87-6.88 (m, 2H), 6.75-6.78 (m, 1H), 3.34-3.35 (m, 2H), 2.91-2.95 (m, 2H), 2.68-2.72 (m, 2H), 1.84-1.95 (m, 4H), 1.45-1.58 (m, 4H), 1.30-1.33 (m, 2H); ^13^C NMR (100 MHz, CDCl_3_) δ 150.3, 131.2, 128.1, 122.5, 113.3, 112.8, 111.9, 103.6, 59.5, 52.9, 50.6, 29.8, 26.3, 24.2, 16.4; Anal. (C_17_H_22_N_2_O · 0.75H_2_O) C, H, N

*2-(2-(1H-indol-3-yl)ethyl)-2-azabicyclo[2.2.2]octane (**4a**).* Compound **4a** was prepared as described for **3a** using 2-azabicyclo[2.2.2]octane (**2**) (222.4 mg, 2 mmol) and 3-(2-bromoethyl)-1H-indole (448.2 mg, 2 mmol) to give the product (370 mg, 73% yield) as a yellow oil. The free base was converted to the HCl salt and recrystallized from methanol to give a tan solid. Mp 251-253 °C; GC/MS (EI) *m/z* 254 (M^+^); ^1^H NMR (400 MHz, CDCl_3_) δ 8.00 (s, 1H), 7.63-7.65 (m, 1H), 7.34-7.36 (m, 1H), 7.04-7.20 (m, 3H), 2.93-2.97 (m, 2H), 2.81-2.89 (m, 4H), 2.66-2.67 (m, 1H), 1.96-2.02 (m, 2H) 1.46-1.72 (m, 7H); ^13^C NMR (100 MHz, CDCl_3_) δ 136.2, 127.6, 121.9, 121.4, 119.1, 119.0, 115.0, 111.0, 57.6, 56.8, 49.6, 25.8, 25.0, 24.4, 24.3; Anal. (C_17_H_22_N_2_ · HCl · 0.25H_2_O) C, H, N.

*2-(3-(1H-indol-3-yl)propyl)-2-azabicyclo[2.2.2]octane (**4b**).* Compound **4b** was prepared as described for **3b** using 2-azabicyclo[2.2.2]octane (**2**) (180 mg, 1.6 mmol) and 3-(1H-indol-3-yl)propanoic acid (302.7 mg, 1.6 mmol) to give the product (350 mg, 82% yield over two steps) as a yellow oil. The free base was converted to the HCl salt and recrystallized from methanol to give a tan foam; GC/MS (EI) *m/z* 268 (M^+^); ^1^H NMR (400 MHz, CDCl_3_) δ 8.00 (s, 1H), 7.61-7.63 (m, 1H), 7.33-7.36 (m, 1H), 7.08-7.20 (m, 2H), 6.98 (m, 1H), ^13^C NMR (100 MHz, CDCl_3_) δ 136.2, 127.6, 121.9, 121.4, 119.1, 119.0, 115.0, 111.0, 57.6, 56.8, 49.6, 25.8, 25.0, 24.4, 24.3; Anal. (C_18_H_24_N_2_ · HCl · 0.5H_2_O) C, H, N.

*3-(2-(8-azabicyclo[3.2.1]octan-8-yl)ethyl)-5-fluoro-1H-indole (**4c**).* Compound **4c** was prepared as for **3b** using 2-azabicyclo[2.2.2]octane (**2**) (180.0 mg, 1.62 mmol) and 3-(5-fluoro-1H-indol-3-yl)propanoic acid (335.4 mg, 1.62 mmol) to give the product (140 mg, 30% yield over two steps) as a yellow oil. GC/MS (EI) *m/z* 286 (M^+^); ^1^H NMR (400 MHz, CDCl_3_) δ 8.51 (s, 1H), 7.20-7.26 (m, 2H), 6.88-7.00 (m, 2H), 2.59-3.67 (m, 7H), 1.90-1.96 (m, 4H) 1.44-1.68 (m, 7H); ^13^C NMR (100 MHz, CDCl_3_) δ 158.8, 156.4, 132.8, 128.0, 127.9, 123.2, 116.4, 116.3, 111.6, 110.2, 109.9, 103.8, 103.6, 62.7, 56.3, 56.1, 49.6, 30.0, 27.9, 25.5, 24.6, 23.9, 22.8; Anal. (C_18_H_23_FN_2_ · 0.5H_2_O) C, H, N.

*Nortropidene (**6**)*. Nortropine (**5**; 10.0 g, 78.6 mmol) was added portion wise to sulfuric acid (10 mL) cooled in an ice bath, then stirred at 160 °C for 4 h. Once cooled to rt, the reaction mixture was diluted with 50 mL H_2_O, 50 mL NaOH (12.5 M), and extracted with Et_2_O (3x 75 mL). The combined organic phases were washed with brine (50 mL), dried over MgSO_4_, and concentrated *in vacuo* to give the title compound as a light orange oil (4.0 g, 47% yield). The free base proved unstable overtime. Thus, it was converted to the HCl salt for long term storage by dissolving in EtOH/conc. HCl, concentrated *in vacuo,* and triturated with DCM/Et_2_O to give a white solid. ^1^H NMR (400 MHz, D_2_O) δ 6.01 (m, 1H), 5.81 (m, 1H), 4.22 (m, 2H), 2.81 (d, *J* = 24 Hz, 1H), 2.27 (m, 3H), 2.13 (m, 1H), 1.96 (m, 1H). HRMS: found m/z = 110.0964 (MH^+^), calcd for C_7_H_12_N (MH^+^).

*1-(8-azabicyclo[3.2.1]oct-2-en-8-yl)-2-(1H-indol-3-yl)ethan-1-one (**7a**).* To a solution of indole-3-acetic acid (1.75 g, 10 mmol) in THF (50 mL) was added CDI (1.95 g, 12 mmol), and the reaction stirred at rt for 2 h. A solution of **6** (1.09 g, 10 mmol) in THF (1 mL) was added to the reaction mixture and stirring continued overnight. The reaction mixture was concentrated, diluted with EtOAc (100 mL), and successively washed with 1N HCl (2x 50 mL), sat. NaHCO_3_ (1x 50 mL), and brine (1x 50 mL). The extract was dried over MgSO_4_, concentrated *in vacuo*, re-dissolved in minimal DCM and precipitated with hexane to give the title compound as an off-white powder and mixture of two diastereomers (A:B, 0.45:0.55) (2.08 g, 78% yield). Diastereomer A: ^1^H NMR (400 MHz, CDCl_3_) δ 8.32 (s, 1H), 7.61 (d, *J* = 8 Hz, 1H), 7.33 (d, *J* = 8 Hz, 1H), 7.12 (m, 3H), 5.99 (m, 1H), 5.47 (m, 1H), 4.89 (m, 1H), 4.33 (m, 1H), 3.79 (m, 2H), 2.43 (d, *J* = 20 Hz, 1H), 2.08 (m, 1H), 1.87 (m, 3H), 1.68 (m, 1H). Diastereomer B: ^1^H NMR (400 MHz, CDCl_3_) δ 8.28 (s, 1H), 7.61 (d, *J* = 8 Hz, 1H), 7.33 (d, *J* = 8 Hz, 1H), 7.12 (m, 3H), 5.81 (m, 1H), 5.47 (m, 1H), 4.79 (m, 1H), 4.33 (m, 1H), 3.79 (m, 2H), 2.82 (d, *J* = 20 Hz, 1H), 2.08 (m, 1H), 1.87 (m, 3H), 1.68 (m, 1H). HRMS: found m/z = 267.1494 (MH^+^), calcd for C_17_H_19_N_2_O (MH^+^).

*1-(8-azabicyclo[3.2.1]oct-2-en-8-yl)-2-(5-fluoro-1H-indol-3-yl)ethan-1-one (**7b**).* To a solution of 5-fluoro-indole-3-acetic acid (0.976 g, 5.05 mmol) in THF (25 mL) was added CDI (0.973 g, 6 mmol), and the reaction stirred at rt for 2 h. The HCl salt of **6** (0.874 g, 6 mmol) was added as a solid, followed by *N,N*-diisopropylethylamine (1.05 mL, 6 mmol) and stirring continued overnight. The reaction mixture was diluted with EtOAc (100 mL), and successively washed with 1N HCl (3x 50 mL), sat. NaHCO_3_ (1x 50 mL), and brine (1x 50 mL). The product was purified by column chromatography (100% DCM to DCM/MeOH/NH_4_OH = 90: 10: 1) to give the title compound as a light peach-colored solid and mixture of two diastereomers (A:B, 0.4:0.6) (1.174 g, 82% yield). Diastereomer A: ^1^H NMR (400 MHz, CDCl_3_) δ 8.16 (s, 1H), 7.25 (m, 2H), 7.14 (d, *J* = 8 Hz, 1H), 6.92 (t, *J* = 8 Hz, 1H), 6.01 (m, 1H), 5.49 (m, 1H), 4.87 (m, 1H), 4.34 (m, 1H), 3.72 (m, 2H), 2.46 (d, *J* = 20 Hz, 1H), 2.12 (m, 1H), 1.90 (m, 3H), 1.72 (m, 1H). Diastereomer B: ^1^H NMR (400 MHz, CDCl_3_) δ 8.16 (s, 1H), 7.25 (m, 2H), 7.14 (d, *J* = 8 Hz, 1H), 6.92 (t, *J* = 8 Hz, 1H), 5.84 (m, 1H), 5.49 (m, 1H), 4.79 (m, 1H), 4.34 (m, 1H), 3.72 (m, 2H), 2.83 (d, *J* = 20 Hz, 1H), 2.12 (m, 1H), 1.90 (m, 3H), 1.72 (m, 1H). HRMS: found m/z = 285.1402 (MH^+^), calcd for C_17_H_18_N_2_OF (MH^+^).

*1-(8-azabicyclo[3.2.1]oct-2-en-8-yl)-2-(5-methoxy-1H-indol-3-yl)ethan-1-one (**7c**).* To a solution of 5-methoxy-indole-3-acetic acid (0.371 g, 1.81 mmol) in THF (10 mL) was added CDI (0.352 g, 2.17 mmol), and the reaction stirred at rt for 2 h. A solution of **6** (0.236 g, 2.17 mmol) in THF (1 mL) was added to the reaction mixture and stirring continued overnight. The reaction mixture was diluted with EtOAc (40 mL), and successively washed with 1N HCl (3x 25 mL), sat. NaHCO_3_ (1x 25 mL), and brine (1x 25 mL). The extract was dried over MgSO_4_, concentrated *in vacuo,* and the residue was purified by column chromatography (100% DCM to DCM/MeOH/NH_4_OH = 90: 10: 1) to give the title compound as an off-white solid and mixture of two diastereomers (A:B, 0.4:0.6) (0.412 g, 77% yield). Diastereomer A: ^1^H NMR (400 MHz, CDCl_3_) δ 8.09 (s, 1H), 7.24 (m, 1H), 7.06 (m, 2H), 6.85 (m 1H), 6.00 (m, 1H), 5.49 (m, 1H), 4.88 (m, 1H), 4.33 (m, 1H), 3.86 (s, 3H), 3.75 (m, 2H), 2.43 (d, *J* = 20 Hz, 1H), 2.11 (m, 1H), 1.88 (m, 3H), 1.66 (m, 1H). Diastereomer B: ^1^H NMR (400 MHz, CDCl_3_) δ 8.06 (s, 1H), 7.24 (m, 1H), 7.06 (m, 2H), 6.85 (m 1H), 5.81 (m, 1H), 5.49 (m, 1H), 4.80 (m, 1H), 4.33 (m, 1H), 3.86 (s, 3H), 3.75 (m, 2H), 2.83 (d, *J* = 20 Hz, 1H), 2.11 (m, 1H), 1.88 (m, 3H), 1.66 (m, 1H). HRMS: found m/z = 297.1601 (MH^+^), calcd for C_18_H_21_N_2_O_2_ (MH^+^).

*(3R,4R,12S,12aR)-2,3,6,11,12,12a-hexahydro-3,12-ethanopyrrolo[1’,2’:1,2]azepino[4,5-b]indol-5(1H)-one (**8a**).* A 25 mL Schlenk tube in an argon atmosphere was charged with **7a** (0.266 g, 1.00 mmol) and Pd(CH_3_CN)_4_(BF_4_)_2_ (0.577 g, 1.30 mmol). Anhyd acetonitrile (10 mL) was added resulting in a dark red solution, which was stirred at rt overnight, maintaining its appearance throughout the time of reaction. The flask was equipped with a deflated balloon, converted to static argon, and a solution of NaBH_4_ (0.113 g, 3 mmol) in 4 mL EtOH was added dropwise over 10 min, precipitating Pd(0) black and filling the balloon with H_2_ gas. The reaction continued to stir for 1 h, where most of the H_2_ was consumed and the product precipitated along with the Pd(0) black. The reaction was diluted with 50 mL 20% MeOH/DCM, vacuum filtered (Pd(0)), and the filtrate successively washed with 1N HCl (1x 20 mL) and brine (1x 20 mL), dried over MgSO_4_ and concentrated *in vacuo*. The product was separated from byproducts by taking up in minimal DCM, diluted with Et_2_O, and filtered to give the title compound as an off-white solid (0.169 g, 63% yield). ^1^H NMR (400 MHz, DMSO-d_6_) δ 10.92 (s, 1H), 7.47 (d, *J* = 8 Hz, 1H), 7.25 (d, *J* = 8 Hz, 1H), 7.03 (t, *J* = 8 Hz, 1H), 6.97 (t, *J* = 8 Hz, 1H), 4.57 (d, *J* = 8 Hz, 1H), 4.27 (d, *J* = 8 Hz, 1H), 3.92 (d, *J* = 16 Hz, 1H), 3.41 (d, *J* = 16 Hz, 1H), 2.98 (d, *J* = 8 Hz, 1H), 2.24 (m, 2H), 2.00 (m, 2H), 1.80 (m, 2H), 1.53 (m, 1H), 1.21 (m, 1H). ^13^C NMR (100 MHz, DMSO-d_6_) δ 171.35, 136.57, 134.78, 127.28, 120.74, 118.39, 117.32, 110.62, 103.88, 55.91, 53.03, 38.29, 32.58, 28.69, 26.92, 25.78, 22.69. HRMS: found m/z = 267.1483 (MH^+^), calcd for C_17_H_19_N_2_O (MH^+^).

*(3R,4R,12S,12aR)-8-fluoro-2,3,6,11,12,12a-hexahydro-3,12-ethanopyrrolo[1’,2’:1,2]azepino[4,5-b]indol-5(1H)-one (**8b**).* Compound **8b** was prepared as described for **8a** using **7b** (0.142 g, 0.50 mmol) and Pd(CH_3_CN)_4_(BF_4_)_2_ (0.289 g, 0.65 mmol) to give the title compound (0.990 g, 70% yield) as a white solid. ^1^H NMR (400 MHz, DMSO-d_6_) δ 11.04 (s, 1H), 7.26 (m, 2H), 6.86 (m, 1H), 4.56 (d, *J* = 8 Hz, 1H), 4.27 (d, *J* = 8 Hz, 1H), 3.90 (d, *J* = 16 Hz, 1H), 3.38 (d, *J* = 16 Hz, 1H), 2.98 (d, *J* = 8 Hz, 1H), 2.23 (m, 2H), 1.99 (m, 2H), 1.79 (m, 2H), 1.51 (m, 1H), 1.18 (m, 1H). ^13^C NMR (100 MHz, DMSO-d_6_) δ 171.19, 156.88 (d, *J*_c_,f = 230 Hz), 138.83, 131.39, 127.61 (d, *J*c,f = 10 Hz), 111.46 (d, *J*c,f = 9 Hz), 108.63, (d, *J*_c_,f = 25 Hz), 104.38, (d, *J*c,f = 4 Hz), 102.34, (d, *J*c,f = 23 Hz), 55.79, 53.02, 38.36, 32.57, 28.64, 26.93, 25.78, 22.70. HRMS: found m/z = 285.1397 (MH^+^), calcd for C_17_H_18_N_2_OF (MH^+^).

*(3R,4R,12S,12aR)-8-methoxy-2,3,6,11,12,12a-hexahydro-3,12-ethanopyrrolo[1’,2’:1,2]azepino[4,5-b]indol-5(1H)-one (**8c**).* Compound **8c** was prepared as for **8a** using **7c** (0.260 g, 0.88 mmol) and Pd(CH_3_CN)_4_(BF_4_)_2_ (0.506 g, 1.14 mmol) to give the title compound (0.210 g, 81% yield) as a white solid. ^1^H NMR (400 MHz, DMSO-d_6_) δ 10.74 (s, 1H), 7.13 (d, *J* = 8 Hz, 1H), 6.99 (s, 1H), 6.67 (d, *J* = 8 Hz, 1H), 4.56 (d, *J* = 8 Hz, 1H), 4.27 (m, 1H), 3.89 (d, *J* = 16 Hz, 1H), 3.76 (s, 3H), 3.41 (d, *J* = 16 Hz, 1H), 2.95 (m, 1H), 2.25 (m, 2H), 2.00 (m, 2H), 1.78 (m, 2H), 1.52 (m, 1H), 1.21 (m, 1H). ^13^C NMR (100 MHz, DMSO-d_6_) δ 171.40, 153.18, 137.25, 129.81, 127.62, 111.26, 110.73, 103.78, 99.48, 55.91, 55.37, 52.99, 38.37, 32.64, 28.69, 26.92, 25.79, 22.78. HRMS: found m/z = 297.1586 (MH^+^), calcd for C_18_H_21_N_2_O_2_: (MH^+^).

*(3R,4R,12S,12aR)-1,2,3,5,6,11,12,12a-octahydro-3,12-ethanopyrrolo[1’,2’:1,2]azepino[4,5 b]indole (**9a**)*. To a suspension of **8a** (0.030 g, 0.11 mmol) in 3 mL THF at rt was added BMS (0.20 mL, 2.1 mmol) and the reaction was stirred at reflux for 1 h. Once cooled to rt, the reaction was slowly quenched with MeOH and concentrated *in vacuo*. The reaction was then taken up in 3 M HCl (5 mL) and stirred at reflux overnight to ensure hydrolysis of the boron complex with the product. The reaction was basified with 1 M NaOH and extracted with DCM (3 x 10 mL). The combined organic phases were washed with brine, dried over MgSO_4_, and concentrated *in vacuo*. The residue was purified by column chromatography (100% DCM to DCM/MeOH/NH_4_OH = 90: 10: 1) to give the title compound as a white solid (0.019 g, 68% yield). ^1^H NMR (400 MHz, CDCl_3_) δ 7.70 (s, 1H), 7.50 (m, 1H), 7.26 (m, 1H), 7.12 (m, 2H), 3.91 (d, *J* = 4 Hz, 1H), 3.49 (m, 1H), 3.38 (m, 2H), 3.17 (m, 1H), 3.03 (m, 1H), 2.80 (d, *J* = 8 Hz, 1H), 2.29 (m, 1H), 2.09 (m, 3H), 1.75 (m, 3H), 1.08 (dd, *J* = 8, 16 Hz, 1H). ^13^C NMR (100 MHz, DMSO-d_6_) δ 139.23, 134.91, 128.27, 120.04, 117.92, 117.17, 111.39, 110.30, 56.25, 56.00, 47.98, 38.39, 29.18, 28.29, 23.88, 23.51, 21.82. HRMS: found m/z = 253.1698 (MH^+^), calcd for C_17_H_21_N_2_ (MH^+^).

*(3R,4R,12S,12aR)-8-fluoro-1,2,3,5,6,11,12,12a-octahydro-3,12-ethanopyrrolo[1’,2’:1,2]azepino[4,5-b]indole (**9b**).* Compound **9b** was prepared as described for **9a** using **8b** (0.028 g, 0.10 mmol) and BMS (0.20 mL, 2.1 mmol) to give the title compound as a white solid (0.019 g, 70% yield). ^1^H NMR (400 MHz, CDCl_3_) δ 7.64 (s, 1H), 7.15 (m, 2H), 6.86 (m, 1H), 3.88 (d, *J* = 8 Hz, 1H), 3.48 (m, 1H), 3.38 (m, 2H), 3.09 (m, 1H), 2.97 (m, 1H), 2.77 (d, *J* = 8 Hz, 1H), 2.29 (m, 1H), 2.10 (m, 3H), 1.74 (m, 3H), 1.09 (dd, *J* = 8, 16 Hz, 1H). ^13^C NMR (100 MHz, DMSO-d_6_) δ 156.74 (d, *J*_c,f_ = 229 Hz), 141.23, 131.56, 128.46 (d, *J*_c,f_ = 10 Hz), 111.77 (d, *J*_c,f_ = 5 Hz), 111.08 (d, *J*_c,f_ = 9 Hz), 107.88 (d, *J*_c,f_ = 25 Hz), 102.11 (d, *J*_c,f_ = 23 Hz), 56.36, 56.22, 47.89, 38.24, 28.85, 27.96, 23.54, 23.20, 21.72. HRMS: found m/z = 271.1604 (MH^+^), calcd for C_17_H_20_N_2_F (MH^+^).

*(3R,4R,12S,12aR)-8-methoxy-1,2,3,5,6,11,12,12a-octahydro-3,12 ethanopyrrolo[1’,2’:1,2]azepino[4,5-b]indole (**9c**).* Compound **9c** was prepared as described for **9a** using **8c** (0.444 g, 1.5 mmol) and BMS (1.50 mL, 15.8 mmol) in THF (40 mL) without the 3 M HCl reflux step to give the title compound as a white solid (0.365 g, 86% yield). ^1^H NMR (400 MHz, CDCl_3_) δ 7.50 (s, 1H), 7.16 (d, *J* = 8 Hz, 1H), 6.96 (s, 1H), 6.79 (d, *J* = 8 Hz, 1H), 3.89 (m, 1H), 3.86 (s, 3H), 3.44 (m, 1H), 3.39 (m, 2H), 3.13 (m, 1H), 2.98 (m, 1H), 2.77 (m, 1H), 2.28 (m, 1H), 2.08 (m, 3H), 1.75 (m, 3H), 1.08 (m, 1H). HRMS: found m/z = 283.1804 (MH^+^), calcd for C_18_H_23_N_2_O (MH^+^).

*(3R,4R,12S,12aR)-1,2,3,5,6,11,12,12a-octahydro-3,12-ethanopyrrolo[1’,2’:1,2]azepino[4,5-b]indol-8-ol (**9d**).* Compound **9d** was prepared as described for **9a** using **8c** (0.450 g, 1.5 mmol) and BMS (1.50 mL, 15.8 mmol) in THF (40 mL) with the 3 M HCl reflux step (12 mL, overnight) to give the title compound as a tan solid (0.078 g, 19% yield). ^1^H NMR (400 MHz, CDCl_3_) δ 7.49 (s, 1H), 7.11 (d, *J* = 8 Hz, 1H), 6.90 (s, 1H), 6.69 (d, *J* = 8 Hz, 1H), 3.88 (d, *J* = 8 Hz, 1H), 3.47 (m, 1H), 3.38 (m, 2H), 3.06 (m, 1H), 2.96 (m, 1H), 2.77 (d, *J* = 8 Hz, 1H), 2.28 (m, 1H), 2.10 (m, 3H), 1.75 (m, 3H), 1.08 (dd, *J* = 8, 16 Hz, 1H). HRMS: found m/z = 269.1654 (MH^+^), calcd for C_17_H_21_N_2_O (MH^+^).

## Supplementary figures

**Supplementary Figure 1.**
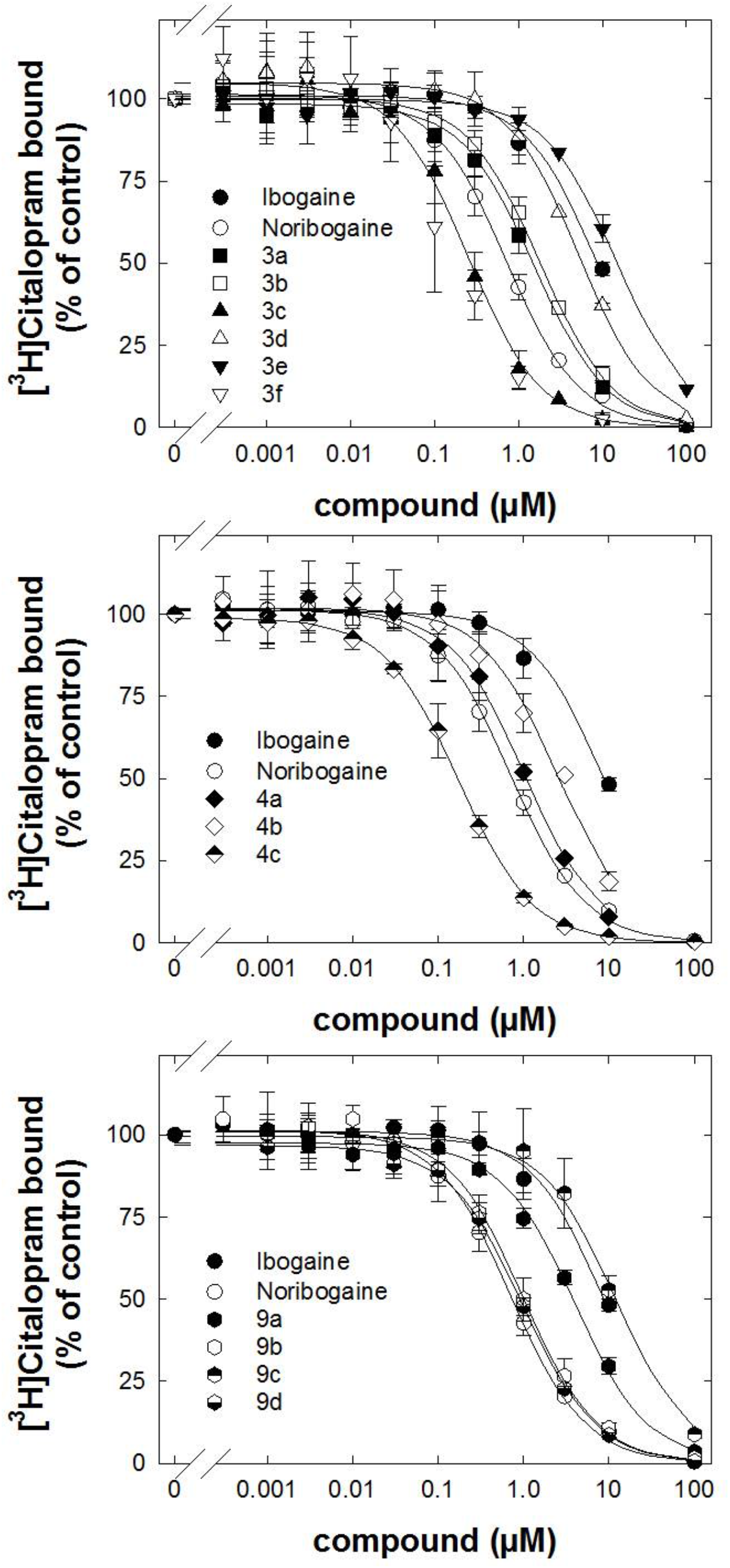
Inhibition by ibogaine analogs of [^3^H]citalopram binding to rat SERT. Brain stem membranes were dissected and prepared from male Sprague-Dawley rat brains (see Methods). Membranes binding was conducted in 96-well polypropylene plates containing 50 μL of various concentrations of the inhibitor, diluted using 30% DMSO vehicle, 300 μL of Tris buffer (SERT), 50 μL of [^3^H]citalopram solution (final concentration 1.5 nM) and 100 μL of tissue (2.0 mg/well original wet weight) for 60 min at room temperature (SERT). Nonspecific binding was determined using 10 μM fluoxetine, which was <10% of specific binding. Data are represented as the means + S.D. (error bars) from at least three independent experiments, each performed in triplicate. Specific binding (between 3000 and 5000 cpm) was set to 100% to normalize for inter-assay variation. The solid curves were drawn by fitting the data to the equation for a monophasic inhibition. *K*_i_-values were calculated from the IC_50_ values using the Cheng-Prusoff equation (see **Table 1)**.

**Supplementary Figure 2.**
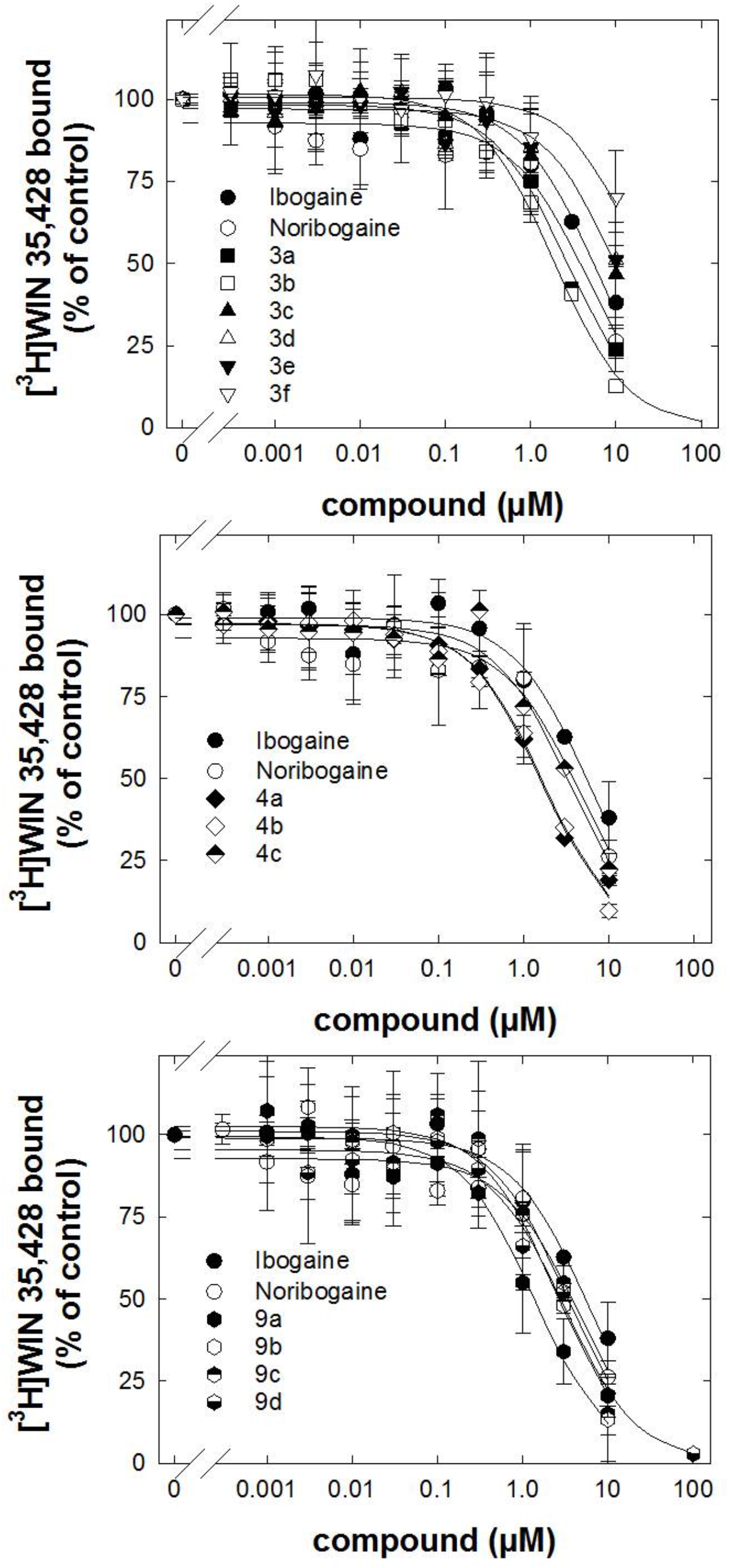
Inhibition by ibogaine analogs of [^3^H]WIN 35,428 binding to rat DAT. Striatum membranes (for DAT) were dissected and prepared from male Sprague-Dawley rat brains (see Methods). Membranes binding was conducted in 96-well polypropylene plates containing 50 μL of various concentrations of the inhibitor, diluted using 30% DMSO vehicle, 300 μL of sucrose phosphate buffer, 50 μL of [^3^H]-WIN35,428 solution (final concentration 1.5 nM) and 100 μL of tissue (2.0 mg/well original wet weight) for 120 min at 0-4 °C. Nonspecific binding was determined using 10 μM indatraline, which was <10% of specific binding. Data are represented as the means + S.D. (error bars) from at least three independent experiments, each performed in triplicate. Specific binding (between 3000 and 5000 cpm) was set to 100% to normalize for inter-assay variation. The solid curves were drawn by fitting the data to the equation for a monophasic inhibition. *K*_i_-values were calculated from the IC_50_ values using the Cheng-Prusoff equation (see Table 1).

**Supplementary Figure 3.**
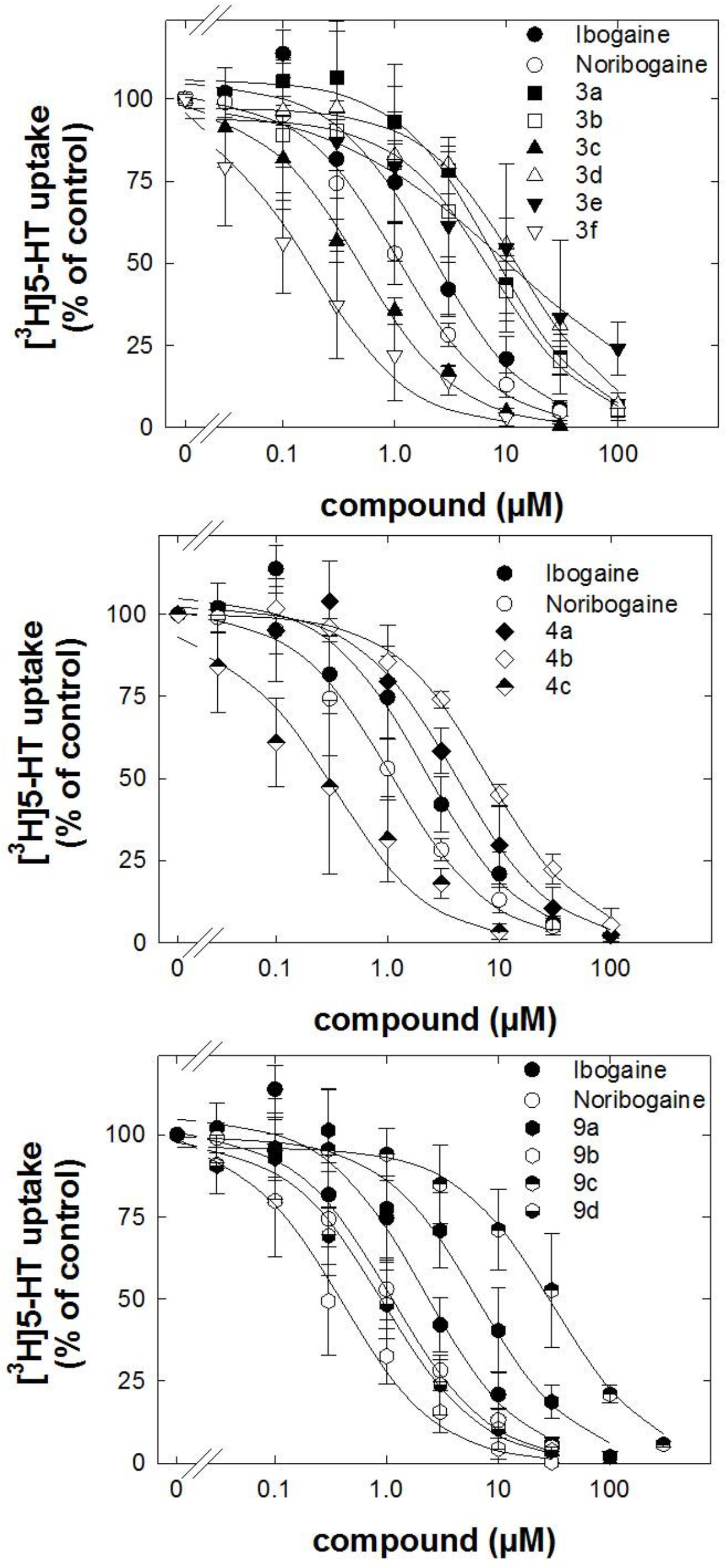
Inhibition by ibogaine analogs of [^3^H]5-HT uptake by hSERT. HEK293 cells stably expressing wild-type YFP-hSERT were seeded onto 96-well plates for 24 h. Cells were incubated with logarithmically spaced concentrations (0.003–300 μM) of ibogaine analogs for 10 minutes and subsequently with the same concentration of the ibogaine analogs with 0.4 μM [^3^H]5-HT for 1 minute. Non-specific uptake was defined as cellular accumulation of radioactivity in the presence of 30 μM paroxetine; this was was <10% of total uptake. Specific uptake is the difference between total and non-specific uptake. Data are the means ± S.D. from three independent experiments done in triplicates. Specific uptake for SERT was 4.46 ± 1.47 pmol·min^-^10^-6^ cells and was set to 100% to normalize for inter-assay variation. The solid curves were drawn by fitting the data to the equation for a monophasic inhibition. The IC_50_-values are reported in **Table 1**.

**Supplementary Figure 4.**
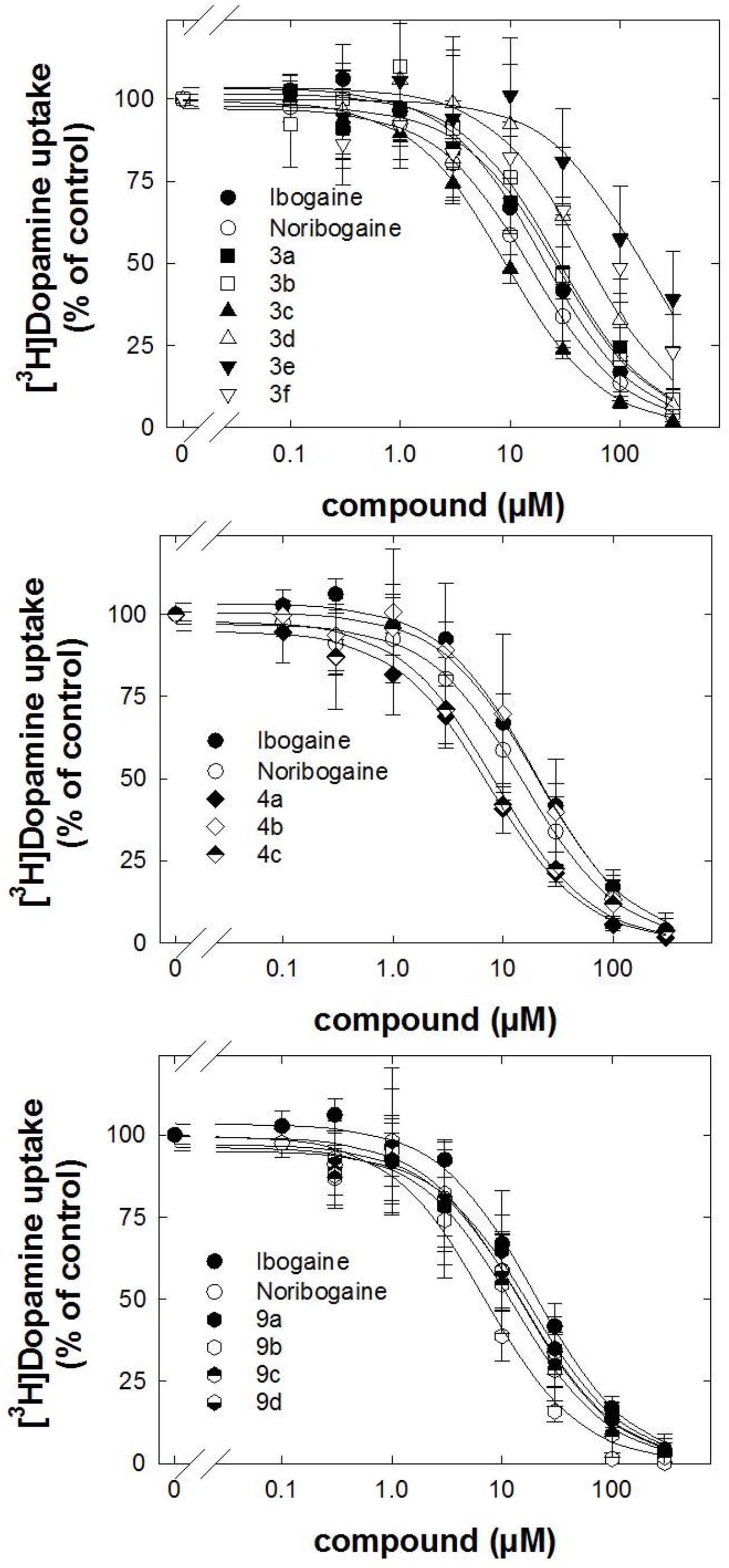
Inhibition by ibogaine analogs of [^3^H]5-DA uptake by hDAT. HEK293 cells stably expressing YFP-hDAT were seeded onto 96-well plates for 24 h. Cells were incubated with logarithmically spaced concentrations (0.003–300 μM) of ibogaine analogs for 10 minutes and subsequently with the same concentration of the ibogaine analogs with 0.4 μM of [^3^H]DA for 1 minute. Non-specific uptake was defined as cellular accumulation of radioactivity in the presence of 30 μM mazindol; this was <10% of total uptake. Specific uptake is the difference between total and non-specific uptake. Data are the means ± S.D. from three independent experiments done in triplicates. Specific uptake for DAT was 7.5 ± 2.1 pmol·min^-^10^-6^ cells, respectively, and was set to 100% to normalize for inter-assay variation. The solid curves were drawn by fitting the data to the equation for a monophasic inhibition. The IC_50_-values are reported in **Table 1**.

